# Genetically modifying the protein matrix of macroscopic living materials to control their structure and rheological properties

**DOI:** 10.1101/2024.05.14.594155

**Authors:** Esther M. Jimenez, Carlson Nguyen, Ahmad Shakeel, Robert Tesoriero, Marimikel Charrier, Alanna Stull, Caroline M. Ajo-Franklin

## Abstract

The field of engineering living materials (ELMs) seeks to engineer cells to form macroscopic materials with tailorable structures and properties. While centimeter-scale ELMs can be grown from *Caulobacter crescentus* engineered to secrete a protein matrix, how the sequence of the protein matrix affects structural and rheological properties remains poorly understood. Here, we explore how changing the elastin-like polypeptide (ELP) length impacts ELM microstructure and viscoelastic behavior. We demonstrate that shortening ELP produces fibers almost 2x thicker than other variants, resulting in a stiffer material at rest. Interestingly, the mid-length ELP forms a complex structure with globules and multidirectional fibers with increased yield stress under flow conditions. Lengthening ELP creates thinner strands between cells with similar storage and loss moduli to the mid-length ELP. This study indicates that sequence-structure-property relationships in these ELMs are complex with few parallels to other biocomposite models. Furthermore, it highlights that fine-tuning genetic sequences can create significant differences in rheological properties, uncovering new design principles of ELMs.

## Introduction

Natural living materials, including bone and biofilms, form complex composites made up of polymers, minerals, and living cells that self-assemble and self-organize into hierarchical structures^1^. These macroscale structures exhibit remarkable characteristics, such as the ability to self-maintain themselves over time, and have tunable rheological properties, which help them thrive in their respective environments^2,3^. Inspired by nature, the field of Engineered Living Materials (ELMs) aims to develop novel materials based on functional combinations of cells and scaffolds^4^. These ELMs have characteristics similar to natural living materials but with tailored functions and properties^5^. So far, the field as a whole has focused on functionalizing ELMs for a variety of applications, including bioremediation and biomedical purposes^6,7^. However, there has yet to be significant work in understanding the sequence-structure-property relationships involving engineered protein matrices in ELMs. This knowledge gap hinders our ability to rationally design protein-based ELMs with desired material properties for real-world applications^8,9^.

One approach to investigating sequence-structure-property relationships in ELMs is to leverage materials designed from the bottom-up. By building new materials from self-assembling cells that create scaffolds, it becomes possible to achieve unprecedented user-defined control over material structure and properties, enhancing material diversity^9,10^. *Caulobacter crescentus* is an attractive platform to develop ELMs from the bottom-up due to its ability to secrete and display its surface layer (S-layer), RsaA, at high levels. RsaA was previously engineered to ligate inorganic, polymeric, and biological materials to the surface of *C. crescentus*^11^, creating a foundation for a 2D assembly system. *C. crescentus* was also modified to secrete non-native biopolymers at unprecedentedly high yields^12^. Building on this work, a macroscopic ELM was developed by engineering *C. crescentus* to secrete and display the Bottom-Up *de novo* (BUD) protein, a self-associating protein that clusters cells and forms an extracellular matrix^13^. The BUD protein consists of RsaA with ELP_60_ replacing the central protein domain. Molinari et al. demonstrated that the rheological properties of the resulting materials called bottom-up de novo ELMs (BUD-ELMs), can be genetically controlled over a factor of 25x through the deletion of large protein domains, including ELP_60_, in the BUD protein. Although broad control over the rheological properties was demonstrated, we have yet to understand how fine-tuned changes to the sequence affect the structural and rheological properties of the resulting material. This limitation prevents ELMs from achieving optimal material performance for a wider range of applications^14^.

In contrast to ELMs, sequence-structure-property relationships in protein-based materials are well established. This paradigm found in Nature has been adopted by material scientists to efficiently design versatile biomaterials composed of proteins such as resilin, collagen, and elastin^8^. These proteins are attractive to study because of their abundance in Nature, multifunctionality, and highly tunable features^15^. For example, elastin-like polypeptides (ELPs), synthetic polymers derived from human tropoelastin, demonstrate a significant range of properties from sequence variations. Its consensus sequence (VPGXG)_n_ can be manipulated by changing position X to any amino acid other than proline. These mutations alter the physical and rheological properties of the resulting ELP-based materials^16,17^. Furthermore, studies done in protein-based block copolymers and recombinant protein elastomers, reveal that changing the tandem repeat length results in different rheological properties^18,19^. Specifically, increasing the tandem repeats in recombinant protein elastomers leads to an increase in tensile strength, a metric describing how much stress the material can withstand before breakage^19^. Although experimental and computational studies have been carried out to understand sequence-structure-property relationships in ELP-based materials, thorough systematic characterization for this paradigm has yet to be done in ELMs hindering predictive material design^20^.

Here, we explore how fine changes to the BUD protein sequence affect the structural and rheological properties of the resulting BUD-ELMs by engineering *C. crescentus* strains expressing BUD proteins with different numbers of ELP tandem repeats. We demonstrate that each strain forms distinct BUD-ELMs with different microstructures varying in fiber directionality and thickness. While all the materials have strong shear thinning behavior, shortening the ELP creates a stiffer material, and the mid-length BUD-ELM has a higher yield stress. Overall, this study provides insight into how small-scale sequence changes lead to different fibrillar thicknesses and orientations, resulting in stiffer and stronger material. The novel sequence-structure-property behavior we observe in our materials sheds light on new conceptual models for ELMs, facilitating material design for future applications.

## Results

### BUD proteins with different ELP repeat lengths are secreted and self-assemble into macroscopic BUD-ELMs

Considerable evidence suggests that increasing the tandem repeats in recombinant ELP elastomers increases the elastic modulus, or stiffness of the material while decreasing the tandem repeats leads to less stiff material^18,19^. To understand if this principle is mimicked in *de novo* living materials, we engineered BUD-ELMs with varying ELP lengths **(Fig. 1A)**. We designed variant Bottom-Up *de novo* (BUD) proteins by replacing the native copy of the surface layer (S-layer) RsaA with a synthetic construct coding four regions using homologous recombination as established in Molinari et al. The first domain in these constructs is the surface anchoring domain of RsaA (*rsaA*_1-250_), allowing monomers to attach to the surface of *C. crescentus*. Following this domain, we inserted a FlagTag, a hydrophilic octapeptide tag, to enable BUD protein detection. Next, we encoded an elastin-like polypeptide (ELP) domain: a synthetic biopolymer derived from tropoelastin composed of a repeating pentapeptide sequence (VPGXG) which self-assembles into a flexible polymer structure in isolation^21^. We chose this biopolymer because it is well-studied and can be recombinantly expressed easily^22^. Following the ELP domain is Spytag, a peptide that forms an irreversible covalent bond to SpyCatcher, its protein partner^23^. Since this split protein system is a well-established covalent functional tag and we know it is functional in RsaA, we used it as a functionalization tag^11^. Lastly, the fourth region encoded is the native secretion sequence of RsaA (*rsaA*_690-1026_), which is known to self-aggregate^24^. These variants were derived from a *C. crescentus* parent strain in which the native S-layer associated protein (*sapA*) is replaced by a xylose-inducible *mKate2*^11^. Each variant has a different tandem repeat length of the ELP: 40 repeats, 60 repeats, or 80 repeats of the VPGXG motif, where X is alanine, glycine, or valine^39^. We refer to these variant BUD protein-expressing strains of *C. crescentus* as BUD_40_, BUD_60,_ and BUD_80_, respectively. The engineered *rsaA* locus of the BUD_40_, BUD_60_, and BUD_80_ strains were sequence verified.

**Figure 1.**
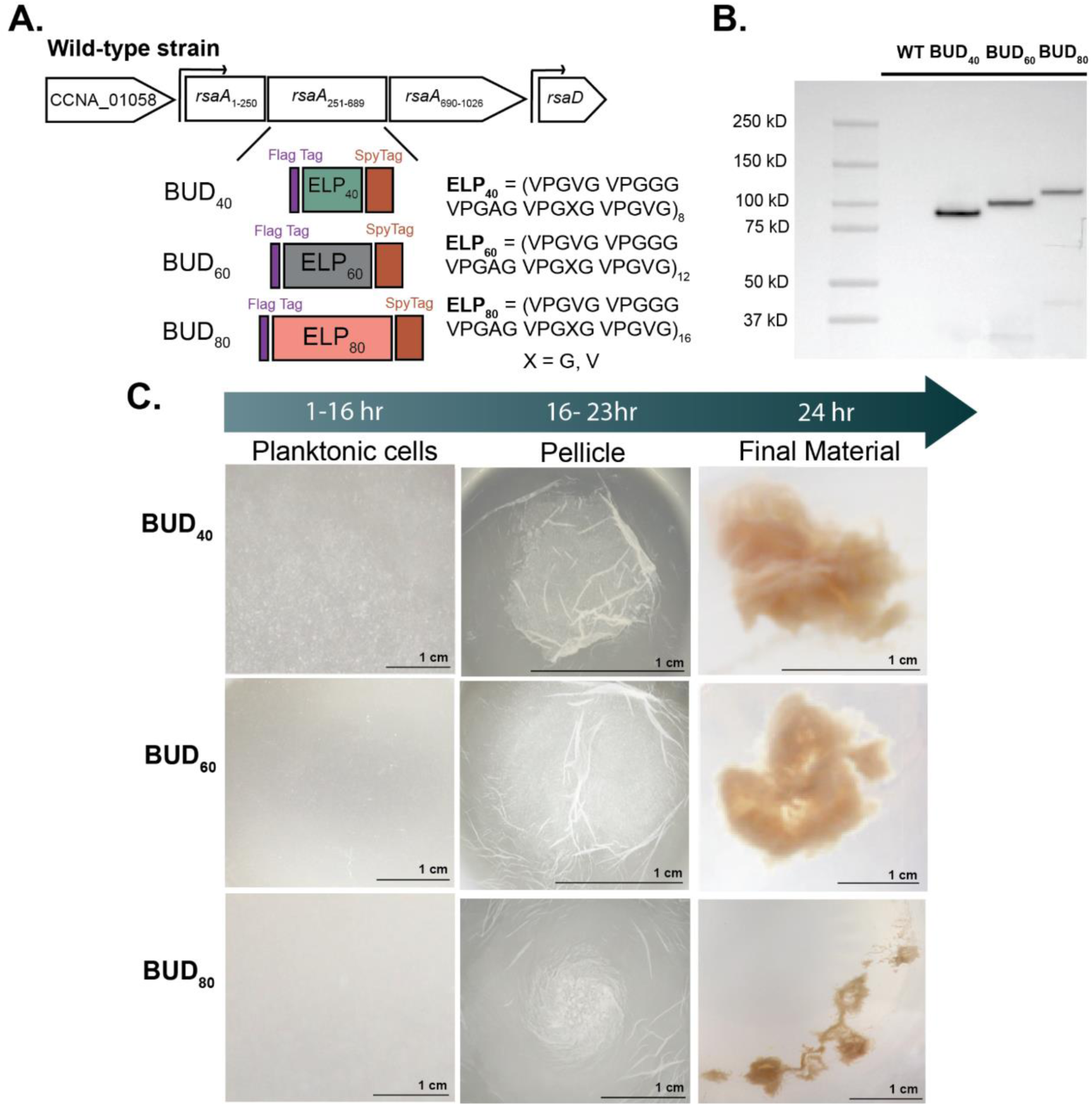
Engineered BUD variant strains assemble into BUD-ELMs. **A)** Schematic of the native *rsaA* gene and the constructs encoding the BUD_40_, BUD_60_, and BUD_80_ proteins consisting of *rsaA*_1-250_, a FlagTag, ELP_n_, Spytag, and *rsaA*_690-1026_. **B)** Immunoblot of culture aliquots using an anti-FlagTag antibody showing BUD proteins are expressed and secreted. Lane 2 is Mfm126 labeled WT for “wild-type”, lane 3 is the BUD_40_ variant, lane 5 is the BUD_60_ variant, and lane 5 is the BUD_80_ variant. The culture from the flasks was harvested after 24 hr in shaking conditions and was not normalized for total protein. C) Optical photographs of the BUD_40_, BUD_60_, and BUD_80_ strain cultures during the planktonic stage (left column), the pellicle stage (middle column), and the fully formed material stage (right column). All variants produce BUD protein and yield a total apparent area of 0.2-2 cm^2^.

To confirm that the full-length BUD proteins are secreted into the extracellular media, we collected aliquots of culture after the cells reached stationary phase and probed these aliquots via immunoblotting with an anti-FlagTag antibody. Prior work shows that the wildtype RsaA and the BUD protein migrate higher than their expected molecular weight^25^. Consistent with their results, the BUD_60_ has an observed band at 100 kDa, despite its expected molecular weight of 98 kDa. The immunoblots illustrate that the BUD_40_ has an observed band at 87 kDa, with its expected molecular weight of 78.5 kDa, and the BUD_80_ has an observed band at 108 kDa while its expected molecular weight was 94 kDa **(Fig. 1B)**. Altogether, this data qualitatively demonstrates that the variant strains can express and secrete BUD proteins with ELPs of different lengths.

To test whether these variant BUD strains form centimeter-scale macroscopic material, we inoculated single colonies in liquid culture using standard media and growth conditions established by Molinari et al. Consistent with prior work, we observed that the BUD_60_ grows cells planktonically for 16 hours before forming a pellicle between 16-18 hours. Following the pellicle stage, the material desorbs from the air-water interface at 24 hours and sinks as the final material. The BUD_40_ and BUD_80_ exhibited slightly different behavior by forming pellicles at a later time point, between 20-23 hours, but both formed material by 24 hours **(Fig. 1C)**. This data indicates that the different biopolymer lengths can still induce material formation via the same assembly steps, although with slightly different time points in the pellicle stage. The differences in assembly did not appear to impact total material production, as image analysis of all BUD-ELM variants yielded a total apparent area of 0.2-2 cm^2^, **(Fig. S1)** suggesting similar sized materials are formed. Interestingly, when removed from the growth flask, the materials appear to retain high amounts of water. To quantify this property, we investigated how much water the materials could hold using thermogravimetric analysis (TGA). These experiments demonstrate that these materials experience a weight loss of 93% **(Fig. S2A)** when heated to 150°C. This indicates the materials hold 93% water by mass and all hold nearly identical amounts of water **(Fig. S2B)**. These results reveal variations in the ELP domain cause modest changes in assembly timing but the size of material produced and level of hydration remain the same.

### Changing the ELP length in the BUD protein produces materials with different fibrillar microstructures

Previous research in ELPs identifies that altering the sequence of the triblock polypeptides by lengthening the hydrophilic domain and incorporating hydrophobic end blocks affects the resulting microstructure of the material^19^. Thus, we hypothesized that varying the hydrophobicity via ELP length impacts the microstructure of the variant BUD-ELMs. To investigate this microstructure, we induced intracellular *mKate2* expression before BUD-ELM formation to visualize the cells, stained the BUD protein matrix after formation using SpyCatcher-GFP, and visualized both components within the BUD-ELMs with fluorescence confocal microscopy **(Fig. 2A)**. Most strikingly, we observed different patterns in the BUD protein-containing matrix between variants. The BUD_40_ contains spider web-like structures that spread between the cells and embed them into the matrix **(Fig. 2B)**. In contrast, the BUD_60_ forms a complex structure consisting of thick and thin fibers surrounded by globular regions of the protein matrix **(Fig. 2C)**. Notably, the BUD_80_ forms thin fiber-like structures surrounded by globular regions of the matrix **(Fig. 2D)**. From the microscopy data, we observed the fibers in the BUD_40_ and BUD_80_ appeared to be more tightly aligned in one direction, while the BUD_60_ appeared to contain randomized fibers with different directionalities. When we quantified the fiber directionalities in the BUD variants using ImageJ (**Fig. S3**), we found that the BUD_60_ has similar amounts of fibers aligned in all directionalities **(Fig. S3B)**. Confirming our visual observations, the BUD_40_ and BUD_80_ had a higher number of fibers aligned over a 20° range **(Fig. S3A, Fig. S3C)**. These findings demonstrate that changing the length of the ELPs in the BUD protein yields changes in the shape and orientation of the protein matrix fibers within the variant BUD-ELMs.

**Figure 2.**
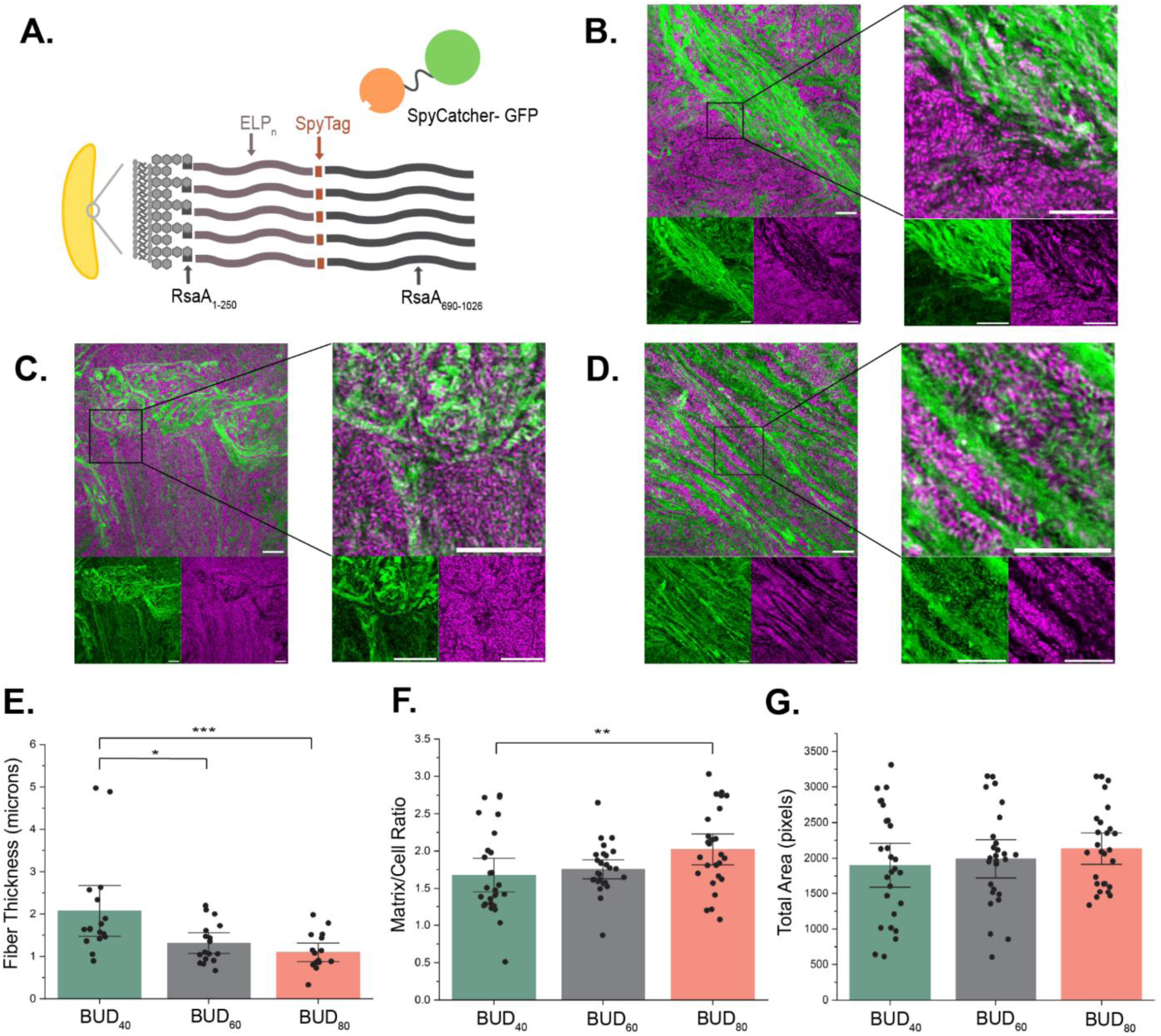
Altering the number of ELP tandem repeats changes the protein matrix microstructure in the BUD-ELMs. **A)** Illustration of the redesigned external surface of *C. crescentus* exhibiting RsaA_1-250_, ELPs of different lengths, SpyTag, and RsaA_690-1026_ attached to the cell’s surface and SpyCatcher-GFP linkage to SpyTag. **B-D)** Confocal microscopy of B) BUD_40_, C) BUD_60_, and D) BUD_80_ materials with xylose-induced cells producing *mKate2* shown in magenta and the BUD protein matrix stained with SpyCatcher-GFP (green). The BUD_40_ and BUD_80_ materials contain aligned fibers of protein matrix, while the BUD 80 protein matrix is a more complex, globular structure. Scale bars are 10 μm for each confocal microscopy figure, and we imaged 3 biological replicates with 3 fields of view per sample. **E)** Comparison of the fiber thickness in microns between the variants showing an increased fiber thickness in the BUD_40_. 17 fibers for each BUD variant were analyzed. **F)** Comparison of the matrix/cell ratio between variants distinguishing significant differences between BUD_40_ and BUD_80_. We analyzed 3 biological replicates with 3 fields of view. **G)** Comparison of the total area in pixels between variants to understand differences in cell clumps. We analyzed 3 biological replicates with 3 fields of view. (*) p-value less than or equal to 0.05, (**) p-value less than or equal to 0.01, and (***) p-value less than or equal to 0.001.

Building upon our understanding of structural differences correlated to varying ELP length, we investigated fiber thickness in the variant BUD protein matrices. To determine if the fibers formed by the matrix vary significantly in width, we analyzed the confocal microscopy images using ImageJ. As a result, we found that the BUD_40_ material produces fibers that are, on average, 2.07± 0.56 microns thick **(Fig. S4)**. In comparison, the BUD_60_ forms fibers about 1.31± 0.23 microns thick, and the BUD_80_ forms fibers that are, on average, 1.07± 0.21 microns thick **(Fig. S5 and Fig. S6)**. Statistical analysis using one-way ANOVA demonstrated a statistically significant difference between the width of the fibers formed by the BUD_40_ and the fibers formed by the BUD_60_ and the BUD_80_ **(Fig. 2E)**. This evidence suggests that shortening the length of the ELP produces thicker fibers within the material.

Having established that changing the ELP length affects the microstructure of the materials, we next sought to understand the composition of the materials. To do so, we analyzed the ratio of matrix to cells in each variant material by splitting the image by fluorescent channel and obtaining the average intensity using ImageJ. Our analysis indicated that differences in the matrix/cell ratio are not statistically significant between the BUD_40_ and the BUD_60_ nor between the BUD_60_ and the BUD_80_ **(Fig. 2F)**. Conversely, the BUD_80_ matrix/cell ratio is significantly higher than the BUD_40_ ratio. To determine if this increase is due to higher levels of BUD_80_ protein secretion, we analyzed the amount of protein produced by each strain with an immunoblot. The results show that the band intensities of both the fraction containing intracellular and cell-bound protein and supernatant fraction for the BUD_40_, BUD_60_, and BUD_80_ are not significantly different. This indicates that both the total amount of protein produced and the amount of protein secreted are similar among all three strains **(Fig. S7A-D).** Thus, lengthening ELP in the BUD protein increases the matrix/cell ratio among the strains but this phenomenon is not due to the protein expression level.

During imaging, we observed several cell-rich regions within all of the variants. Therefore, to establish if the variants differ in overall cell content, we quantified the total cell area present in the confocal microscopy images using ImageJ **(Fig. 2G)**. To achieve this goal, we segmented the images to create binary masks of the cell content in each image, from which we measured the total cell area **(Fig. S8)**. We found that the total area of cell-rich regions is not statistically significant between the variant materials. This indicates that the amount of cell content within these materials does not differ.

### The variant BUD-ELMs have different stiffness and yield stress behavior

Having observed differences between the material microstructures, we sought to investigate if these changes affect how the materials respond under deformation forces, also known as their rheological behavior^26^. Rheological tests enable a material’s storage and loss moduli, viscosity, and yield stress to be measured^27^. Previous research indicates that decreasing the tandem repeats in purified elastin-like polypeptides leads to stiffer properties, while increasing the number of tandem repeats leads to more flexibility^18,19^. We hypothesized that this characteristic would be imitated in the BUD-ELMs.

To characterize the rheological behavior of the variant BUD-ELMs, we measured their rheological properties. To facilitate the integrity of the cells in the material, the material should remain hydrated throughout the experiment. Therefore, we conducted standard oscillatory time sweeps on the BUD_60_ to establish the time point at which these materials start to change rheological properties due to dehydration. The results revealed that the materials storage modulus started increasing after 20 minutes on the rheometer, indicating the material started dehydrating **(Fig. S9),** and therefore, future tests were carried out within 20 minutes. Because the BUD_60_ held similar amounts of water as the BUD_40_ and BUD_80_, we assumed the dehydration rate would be similar and completed the experiments within 20 min for all samples.

To determine the linear viscoelastic region (LVE) and the crossover point, we performed oscillatory amplitude sweeps. The LVE indicates the oscillation strain range in which frequency sweep tests can be carried out without breaking the polymer network. The crossover point is the oscillation strain (%) where the material transitions from solid-like to liquid-like behavior, represented by storage modulus (G’) and loss modulus (G”), respectively. Amplitude sweeps illustrated that the LVE for each variant was between 0.1%-1% oscillation strain **(Fig. 3A)**. Before the 1% oscillation strain, the materials were a viscoelastic solid. However, after this threshold, the G’ began to decrease, and the materials began to lose their structural integrity, displaying viscoelastic liquid behavior. On an average of 10 replicates, the BUD_60_ has a crossover point of 10.9% ± 0.74, while the BUD_40_ and BUD_80_ have lower crossover points, 8.16% ± 0.94, and 8.14% ± 0.94, respectively **(Fig. 3B)**. Together, these results demonstrate a higher (%) strain is necessary for the BUD_60_ to flow and exhibit viscoelastic liquid behavior.

**Figure 3.**
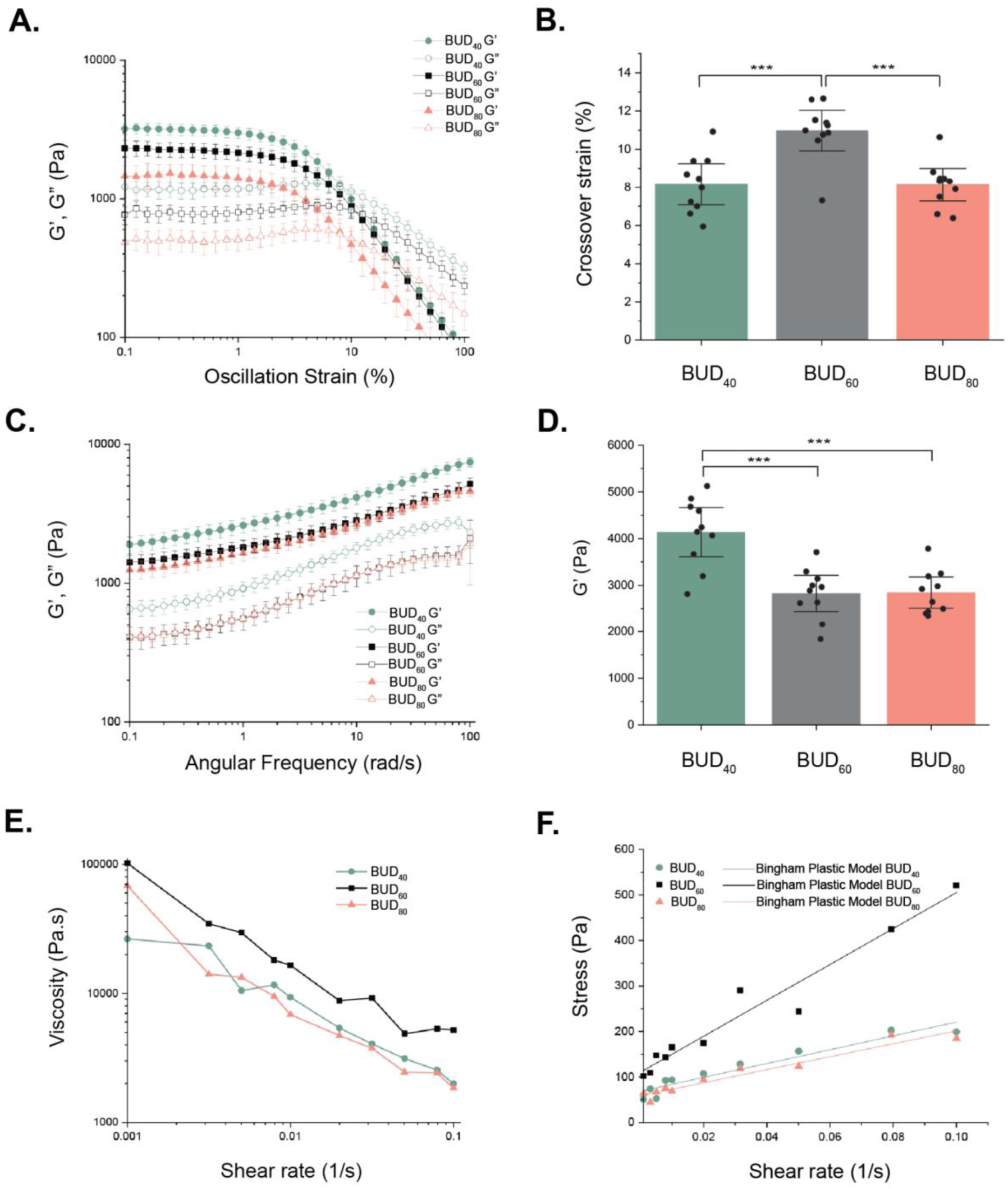
Variant BUD-ELMs have different rheological properties. **A)** Oscillation strain sweeps were acquired from 0.1% to 100% strain amplitude at a constant frequency of 3.14 rad/s identifying the linear viscoelastic region of 0.1-1% for the materials. The storage (G’) and loss (G”) moduli for BUD_40_, BUD_60,_ and BUD_80_ were plotted. **B)** The crossover strain (%) of each BUD material was collected from the crossover between G’ and G”. The BUD_60_ has a higher crossover strain (%) indicating greater resistance to deformation. **C)** Frequency sweep measurements were acquired from 0.1 to 100 rad/s at a constant strain amplitude of 1%, demonstrating the BUD_40_ produces a stiffer material. D) The average storage (G’) modulus of the BUD_40_, BUD_60_, and BUD_80_ at an angular frequency of 10 rad/s showing the differences between the stiffness of the materials. Error bars are centered on the mean value and represent 95% confidence intervals of 10 samples. **E)** The viscosity vs. shear rate of the variants exhibiting strong shear-thinning. **F)** Stress vs. shear rate of the variants fitted using the Bingham plastic model to compare yield stress values at the y-intercept. The mean value of 2 samples is plotted. (*) p-value less than or equal to 0.05, (**) p-value less than or equal to 0.01, and (***) p-value less than or equal to 0.001.

As seen in other ELP-based protein systems, we hypothesized that the BUD_40_ strain would produce a stiffer material because shorter ELP tandem repeats tend to form stiffer materials. To probe G’ and G’’, we performed frequency sweeps at a 1% oscillation strain on all three materials. Rheological measurements confirmed that all three variants were viscoelastic solids since G’ was significantly higher than G” (**Fig. 3C**). Additionally, their storage moduli ranged between 1,000-10,000 Pa. As expected, the frequency curve identified a significantly higher G’ and G” for the BUD_40_ compared to the other variants throughout the angular frequency range of 0.1-100 rad/s **(Fig. 3C)**. The BUD_60_ and BUD_80_ have almost identical G’ and G” values, indicating similar rheological footprint behavior. To present a quantitative comparison between all the materials’ G’, we chose values at an angular frequency of 10 rad/s. The results showed that the BUD_40_ G’ increases by 1,000 Pa compared to the BUD_60_ and BUD_80_ **(Fig. 3D)**. This data suggests that decreasing the tandem repeats of ELP increases the material’s stiffness. Overall, this test demonstrates that through genetic modification of the BUD protein, the change in stiffness of the BUD-ELMs mimics qualitative trends seen in purified ELPs.

Hydrogels are shear-thinning materials because their weak polymer networks break relatively easily under shear stress^28^. Given the similarities between BUD-ELMs and hydrogels, we hypothesized these variant BUD-ELMs might be shear-thinning materials^29^. To assess this, we employed an equilibrium flow test by applying a constant shear rate until the stress reached equilibrium and analyzed the relationship between viscosity and shear rate. We observed that the viscosity decreased as the shear rate increased for all three BUD-ELM variants **(Fig. 3E)**. Furthermore, when we fitted the data to the Power Law model, we found that the power law index is similar for all BUD variants **(Fig. S10)**. This trend indicates that all the materials show shear-thinning behavior but that changing the number of tandem repeats in ELP does not affect the shear-thinning behavior of the materials.

Next, we wanted to determine the yield stress - the minimum force required to break the material’s microstructure at rest to make it flow^30^ - of the BUD-ELM variants. To obtain the yield stress of these materials, we measured the stress as a function of shear rate and fit the resulting trends to the Bingham plastic model^31^ **(Fig. 3F)**,

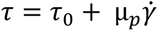

where *τ* is the shear stress, τ_0_ is the yield point, *µ_p_* is the Bingham viscosity, and γ is the shear rate. The trends were well-described by this linear relationship, with R^2^ for the BUD_40_, BUD_60_, and BUD_80_ of 0.92, 0.95, and 0.94, respectively. Because the crossover strain (%) of the BUD_60_ variant was higher than the other variants **(Fig 3B)**, we hypothesized that the BUD_60_ would have a higher yield stress. We found that the yield stress values for the BUD_40_ and BUD_80_ are very similar, 68.80 ± 7.05 Pa and 59.11 ± 5.69 Pa, respectively **(Fig. 4A)**. The yield stress of the BUD_60_ is 110.48 ± 14.1 Pa, which is almost two times higher than the other materials. The Bingham viscosity follows the same trend as the yield stress, showing the viscosity of the BUD_60_ at high shear rate is higher than the other variants **(Fig. 4B)**. Altogether, this data indicates that under flow conditions, the BUD_60_ exhibits higher yield stress and Bingham viscosity, indicating these parameters are non-linearly related to the number of ELP tandem repeats.

**Figure 4.**
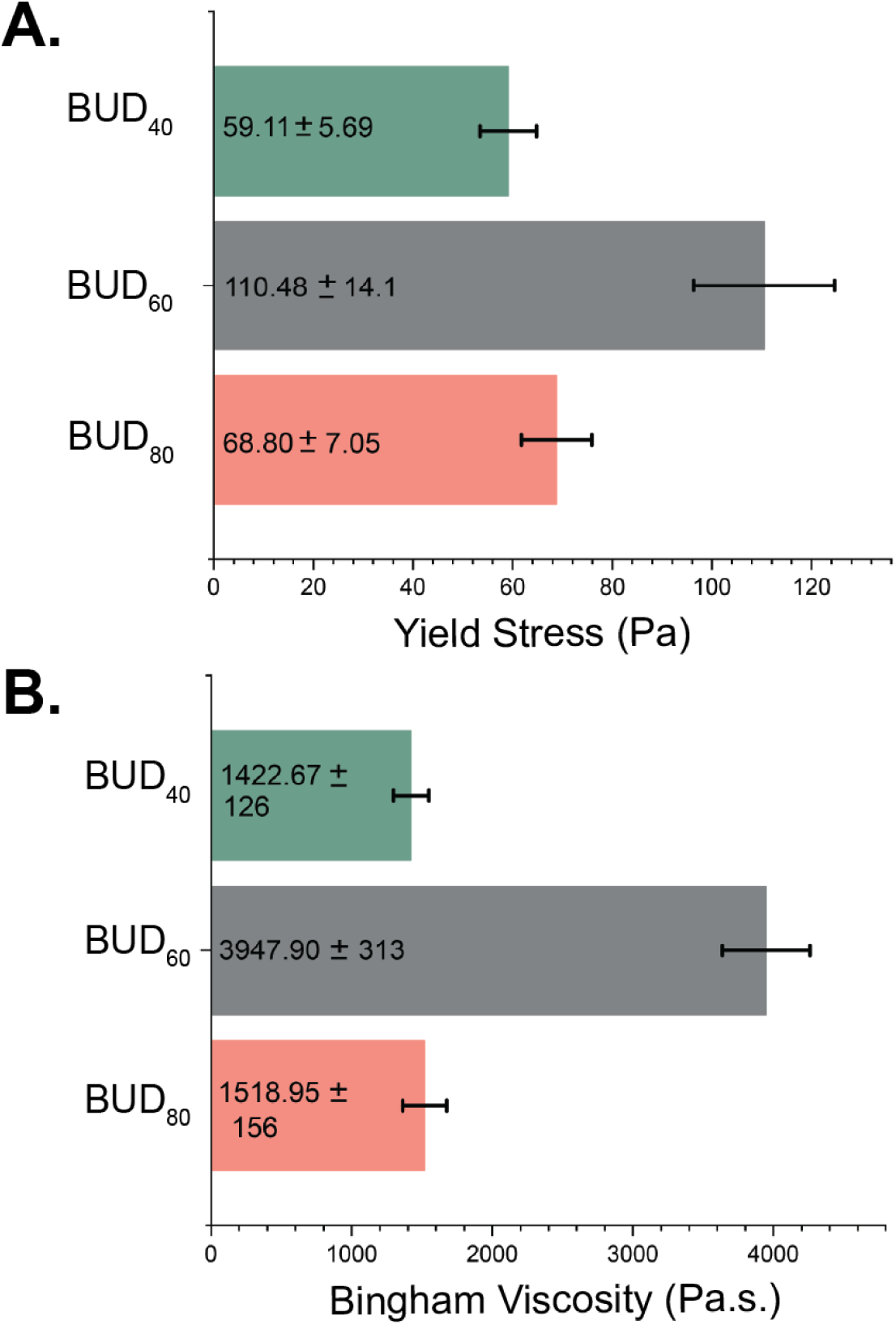
Yield Stress and Bingham Viscosity for the BUD variants. **A)** Average yield stress values for the BUD variants. Consistent with the amplitude sweeps, the BUD_60_ has a higher yield stress. **B)** The mean Bingham viscosity for the variants showing higher average values for BUD_60_. Error bars are centered on the mean value and represent the standard error of 2 technical samples.

## Discussion

In summary, our results show that altering the ELP length in BUD-ELM drives complex, multifaceted changes in their microstructure and rheological properties. Encoding different length ELPs in the BUD protein produces centimeter-scale variant BUD-ELMs. Modest alterations in the length of the ELP in the BUD protein vary the orientation and thickness of fibers in the resulting material and their rheological properties. The thicker fibers of BUD_40_ are correlated with a stiffer material. The randomized complex network of fibers in the BUD_60_ is associated with a 2x higher yield stress and plastic viscosity compared to the BUD_40_ and BUD_80_ variants. This work reveals connections between genetic sequence, fiber morphology, and rheological properties in ELMs that begin to elucidate sequence-structure-property in these emerging materials.

This study suggests that microstructure characteristics are a function of ELP sequence length in BUD-ELMs. All the variant BUD-ELMs formed fibers in their microstructure. This is consistent with previous work where researchers show that “double-hydrophobic” ELP block copolymers made up of the sequence, (VGGVG)_5_-(VPGXG)_25_-(VGGVG)_5_ generate nanofibers^32^. Surprisingly, we also see the thickness of the microfibers are inversely correlated with the length of the ELP: longer ELPs generate thinner fibers. We put forth two possible explanations for this trend. First, the longer ELPs could act as more flexible linkers, decreasing the nucleation of individual fibers and thus decreasing their width. A similar phenomenon was observed with silk fibers that nucleated on the surface of *B. subtilis*: silk peptides with less flexible linkers increased nucleation leading to thicker fibers^33^. An alternative explanation is that the decreasing fiber thickness arises from increasing hydrophobicity of the longer ELPs. The sequence of our ELP means that the overall hydrophobicity of the ELP increases as the tandem repeat length increases^34^. Thus, the difference in hydrophobicity between these variants may play a role in the fiber formation process. More broadly, the patterns in fiber microstructure observed in this work cannot be explained by previously reported sequence-structure relationships for existing biomaterials, such as biofilms, cellulose, or silk. This further demonstrates the need for in-depth characterization of ELMs to develop models. Additional studies are underway to examine the BUD protein assembly process and identify the interactions responsible for the differences in fiber formation.

Our work also suggests that specific microstructure characteristics are connected to certain rheological properties within BUD-ELMs **(Fig 5)**. We observed that the BUD_40_ produces stiffer materials at rest compared to the BUD_60_ and BUD_80_. We suggest this may arise from the thick fibers produced by the matrix. In addition, we found that the BUD_40_ and BUD_80_ have lower yield stress and plastic viscosity than the BUD_60_. Hypothetically this is due to the oscillation shear stress being imposed in a direction parallel to the fiber alignment (fibers aligned in the direction of shear), causing the strength exerted by the fibers to be lower than if the stress was imposed in the perpendicular direction as seen in cellulose nanofibers^35^. We do not know the direction in which the fibers are aligned in the material during rheometry; therefore, we cannot control the direction of the stress with the fibers. In contrast, the BUD_60_ has a complex, randomized microstructure, where the fibers are aligned in various directions, achieving resistance to flow in different directions and causing the material to be high strength^36^.

**Figure 5.**
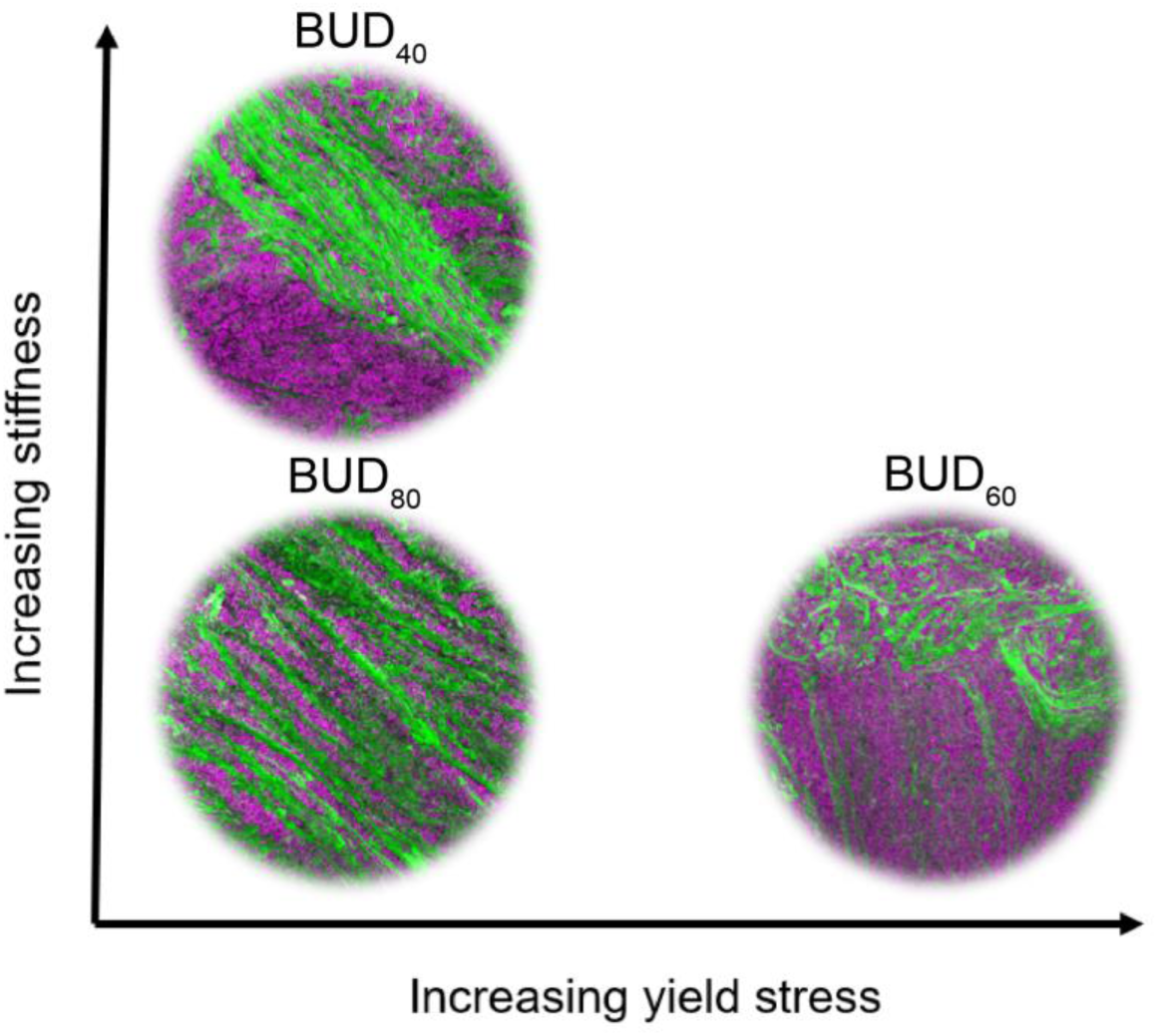
Illustration connecting the sequence-structure-property relationships of the BUD variants. The BUD_40_ produces thicker fibers that contribute to an increased stiffness of the material at resting conditions. Under flow conditions, the BUD_60_ creates a complex structure consisting of matrix globules and non-directional fibers that help increase yield stress.

Taken together, this study advances our insight into the sequence-structure-property relationships in three ways. The first insight is an increased understanding of the rheological behavior of these BUD-ELMs. This work enabled us to identify these BUD-ELMs as shear-thinning materials that could be applied to biomedical applications, such as drug delivery and 3D printing^37,38^. The second advancement is that we established methods to comprehensively characterize the macroscopic, microscopic, and rheological properties of these biomaterials. Third, we obtained insight into how changes to the primary sequence of these proteins create drastic changes in the properties of the materials.

With these insights in hand, forthcoming research should focus on uncovering the diversity of sequences that can self-assemble into ELMs and the relationship between biopolymer sequence length, fiber thickness, and fiber alignment. We envision creating a database of biopolymers that can be swapped into BUD-ELMs to target desired properties such as durability and elasticity for applications in energy, environment, and medicine. Additionally, because this work emphasized that small-scale sequence changes can fine-tune the structural and rheological properties of *de novo* biomaterials, we see future work is needed to develop new conceptual models for resulting emergent properties in BUD-ELMs.

## Conclusion

Learning to genetically manipulate ELMs for the predictive design of materials will allow for reduced time spent testing many variants to achieve necessary rheological properties and greater efforts in functionalizing ELMs. This is the first comprehensive study demonstrating the importance of sequence-structure-property relationships in ELMs. This work shows that in BUD-ELMs, a randomized fiber network allows for resistant behavior in flow conditions, while thick fibers help stiffen the material in rest conditions. Thick fibers producing stiffer materials have never been noted in biomaterials and this phenomenon shows the importance of fundamental characterization of ELMs. We anticipate this insight will have applications to 3D printing new living devices, drug delivery, and tissue engineering. The new design principles for BUD-ELMs herein opens a new avenue of biomaterials research that will revolutionize predictive design.

## Acknowledgments

We gratefully acknowledge the financial support of the National Science Foundation Graduate Research Fellowship Esther M. Jimenez (ID 2022300506), CPRIT to Caroline M. Ajo-Franklin. (award # RR1900063), and the Welch Foundation to Caroline M. Ajo-Franklin (award # C-2130-20230405). We thank our colleagues Dr. Leanne M. Friedrich and Swetha Sridhar for fruitful scientific conversations.

## Methods

### Construction of variant BUD-ELM strains

To generate the variant BUD-ELM strains (BUD_40_-gCAF018 and BUD_80_-gCAF019) from the wild-type (MFM126), we modified the pNPTS138-based integration vector pSMCAF008, which was used to build the original BUD-ELM strain established by Molinari et al. 2022. By replacing the ELP_60_ sequence from pSMCAF008 with two in-frame BsaI sites, we created a universal backbone for BUD-ELM constructs (pBTCAF008). The sequences for *ELP_40_, ELP_80_,* and S*pyTag* were then inserted into this backbone by Golden Gate Assembly. All ELP sequences were obtained from GenScript, and the SpyTag sequence was obtained from Twist Bioscience. The full sequences are found in the supplementary information.

To generate the variant BUD-ELM strains, the plasmids pCAF216 (BUD_40_) and pCAF215 (BUD_80_) were first electroporated into *C. crescentus NA1000 ΔsapA::*Pxyl-mKate2 (MFM126). The culture was then plated on PYE with 25 μg/ml kanamycin to select for integration of the plasmid. Successful integrants were grown overnight at 30°C in 2 ml liquid PYE without selection to allow for recombination. Then 50 μL of each culture was plated on PYE + 3% sucrose to select against cells possessing the *sacB* gene, allowing for successful isolation of cells with the correct BUD-ELM genotype. Successful integration of the sequences was confirmed using touchdown PCR with annealing temperature ranging from 72°C-62°C, decreasing 1°C per cycle, and the PCR amplifications were sequenced using GENEWIZ. All the strains and plasmids generated in this study are found in Table S1 and Table S2.

### Growth conditions of BUD-ELMs

The BUD-ELMs were grown by inoculating a colony of the *C. crescentus* strains into 80 ml of PYE in a 250 ml glass flask. The cultures were grown in an incubator at 30°C at 250 rpm with a 2 inch-orbit for 24 hrs.

### Immunoblot analysis of BUD proteins

Cultures of *C. crescentus* of BUD-ELM strains (gCAF018, RCC002, and gCAF019) were grown in shaking conditions until they reached the stationary phase. 1 ml supernatant was taken from each culture flask and diluted with 2x Laemmli buffer, which was loaded onto a TGX Stain Free gel (Bio-Rad Laboratories). After running, the gel was transferred to a 0.2 μM nitrocellulose membrane, left on the bench to dry for 30 min, and then blocked for 1 hr at room temperature with SuperBlock blocking buffer (Thermo Scientific). Membranes were then washed five times in TBST buffer and then incubated in a 1:5000 dilution of Monoclonal ANTI-FLAG antibody for 1 hr at room temperature. Membranes were washed an additional five times in Tris Buffered Saline with 0.1% Tween (TBST) buffer. Then, Clarity Max Western ECL Substrate (Bio-Rad Laboratories) was added to the membrane and imaged immediately for chemiluminescence. Image analysis was performed using the AlphaView software installed in the FluorChem E system.

### Immunoblot analysis for Protein secretion analysis

Cultures of *C. crescentus* of BUD-ELM strains (gCAF018, RCC002, and gCAF019) were grown in shaking conditions until they reached 21 hrs. The OD was taken for each culture and normalized to 1 OD. To separate the pellet from the supernatant, the culture was collected into a 15 ml tube and spun for 15 min at 4000 xg using the Avanti J-15 R tabletop centrifuge (Beckman Coulter). Then, to concentrate the supernatant until it was 500 μl, the Amicon Ultra-0.5 ml centrifugal filters (Milliporesigma) were used and spun for 10 min at 14,000 g on the Eppendorf 5424 microcentrifuge. The pellet was resuspended with 1 ml PBS. 10 μl of the resuspended pellet and concentrated supernatant for each culture was diluted with 2x Laemmli buffer, which was loaded onto a TGX Stain Free gel (Bio-Rad Laboratories). After running, the gel was transferred to a 0.2 μM nitrocellulose membrane, left on the bench to dry for 30 min, and then blocked for 1 hr at room temperature with SuperBlock blocking buffer (Thermo Scientific). Membranes were then washed five times in TBST buffer and then incubated in a 1:5000 dilution of Monoclonal ANTI-FLAG antibody for 1 hr at room temperature. Membranes were washed an additional five times in Tris Buffered Saline with 0.1% Tween (TBST) buffer. Then, Clarity Max Western ECL Substrate (Bio-Rad Laboratories) was added to the membrane and imaged immediately for chemiluminescence. Image analysis was performed using the AlphaView software installed in the FluorChem E system.

### Apparent BUD-ELM size measure

To elucidate potential size differences between BUD-ELM strains, flasks of each strain were grown under standard conditions before the bottom of each flask was imaged using a Canon EOS 77D camera. Flasks were positioned within a reflective photobox on a clear plastic surface such that the bottom of the flask stood approximately 11.5 cm above the camera lens. These images were separated into RGB channels using MATLAB R2023b. A subset of the blue channel images were then input into the image classification software ilastik (version 1.3.3), as a training set for the autocontext workflow. The first stage of training separated images into three different classifications: background, Artefact and Sheared Material, and Material. Artifacts and Small Material were defined as remaining scratches and stains on the flask that might impact analysis as well as small, sheared pieces of material that would lead to systematic overestimations of material area. The Material group therefore referred to larger, connected pieces of material. The second stage of training distinguished these larger pieces of material from the rest of the image. Both stages of training utilized all 37 features provided within the ilastik workflow (Ilastik, version 1.3.3). From the results of the training set, the second stage segmentation masks for all blue channel images were calculated, and loaded into MATLAB R2023b. From these masks, the flat area of each piece of material was calculated; to eliminate any remaining artifacts from analysis only objects possessing an area equal to or greater than 5% of the total area of material for that given flask were utilized. Additionally, for each image, a conversion rate between pixels and squared centimeters was determined using the standard flask diameter as a reference point. After conversion, the total area of material per flask, the largest piece of material per flask, and the distribution of material sizes were measured.

### Thermogravimetric Analysis of BUD-ELMs

Variant BUD-ELMs (gCAF018, RCC002, and gCAF019) were grown in 80 ml PYE at 30°C at a shaking speed of 250 rpm for 24 hrs. The samples were collected and weighed around 13.5 mg. They were heated in a high-temperature 70 μL Aluminum pan from 25^°^C - 150^°^C at a ramp rate of 5°C/min. TGA experiments were performed using a Mettler Toledo TGA/DSC 3+ STARe System.

### Confocal Microscopy

Single colonies of variant BUD-ELM strains (gCAF018, RCC002, and gCAF019) were inoculated in 80 ml PYE with 0.15% D-xylose- to induce the expression of *mKate2*, in a 250 ml flask and grown in 24 hrs at 30°C at a shaking speed of 250 rpm. The material was collected and washed twice with 1 ml of 0.01 M Phosphate-buffered saline (PBS), in an Eppendorf tube. Then they were incubated with 80 μg/ml of SpyCatcher-GFP for 1 hr. After incubation, the samples were washed three times with 1 ml 0.01 M PBS and then some of the material was placed between a glass coverslip-bottomed 50 mm Petri dish with a glass diameter of 30 mm (MatTek Corporation) and a slab of PYE agarose (1.5% w/v). The Zeiss LSM800 Airyscan confocal microscope was used for imaging. Data was analyzed using ImageJ software.

### Matrix/Cell Ratio Analysis

The cell fluorescence intensity for the cells and matrix were analyzed using ImageJ. The intensity of 27 images was averaged for the cells and matrix. The matrix/cell ratio was calculated by dividing the mean intensity for the matrix by the mean intensity for the cells.

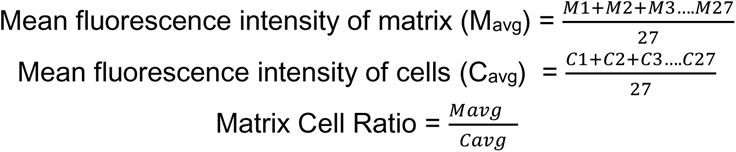

### Cell clump Analysis

We assumed that the more black pixels exposed in the image, the more cell clumps are within the image, and the fewer black pixels in the final image, the fewer cell clumps in each image. To obtain the total black pixeled area that did not have cells, the threshold was adjusted to highlight the cell clumps in black. Then, a binary mask of this black-and-white image was created to highlight the area that does not have cells in black. The black area was then measured and an average was taken using the 27 samples.

### Rheological measurements

The variant BUD-ELMs were grown in standard conditions and then collected into a 1.5 ml Eppendorf tube and spun for 10 s at 3200 rcf using a mini centrifuge (VWR - C0803). Then the supernatant was removed, and 200 μL of fresh PYE was added to the BUD-ELMs to prevent the material from getting dry. The rheological properties of the variant BUD-ELMs (gCAF018, RCC002, and gCAF019) were evaluated on a strain-controlled ARES G2 rheometer with a 0.1 rad 8-mm diameter cone plate geometry. Preshearing was performed before each test at a 1% shear rate for 30 s with a 10 s equilibration time. Small amplitude time sweeps were performed for 3600 s at a 0.5% oscillation strain and a fixed frequency of 0.1 rad/s. Strain sweep experiments from 0.1 to 100% oscillation strains were performed at a fixed frequency of 3.14 rad/s. Frequency sweep experiments were conducted at a 1% strain amplitude from 100 to 0.1 rad/s. Equilibrium stress growth tests were performed at constant shear rates of 0.001, 0.00316, 0.00501, 0.00794, 0.01, 0.01996, 0.03163, 0.05013, 0.07944, 0.1 for each variant until stress equilibrium is reached with a 2% tolerance within 3 consecutive points. This data was further used to get the flow curve which was plotted as viscosity (Pa.s) vs. shear rate (1/s) and shear stress (Pa) vs. shear rate (1/s). Data were acquired using TRIOS software (version 4.2.1.36612) and analyzed using Origin (version 9.9.0.225).

