## Supplemental Information for "Genetically modifying the protein matrix of macroscopic living materials to control their structure and rheological properties"

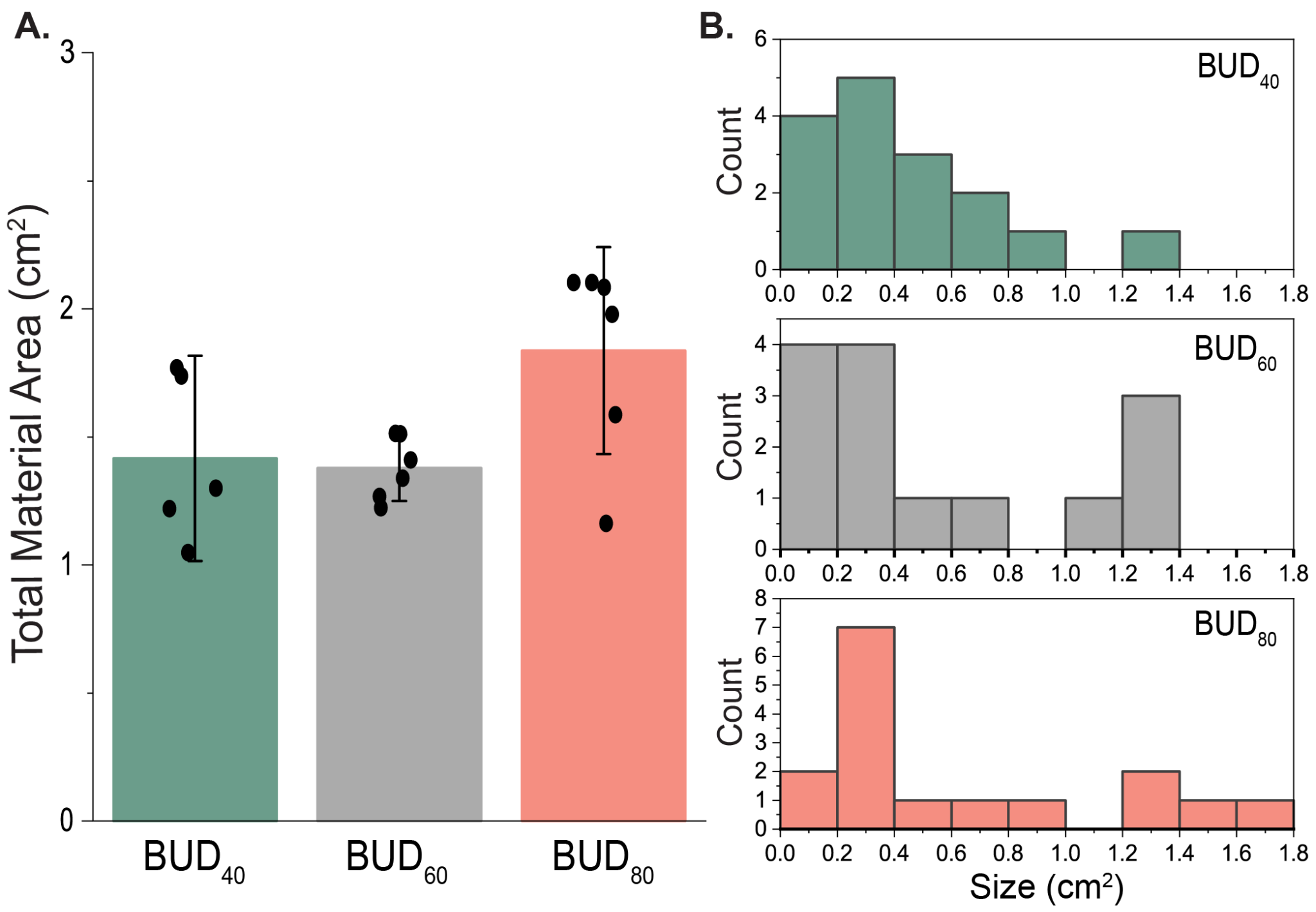
**Figure S1.** **The total material area remains the same for the variant BUD-ELMs. A)** Total Material Area of the variant BUD-ELMs produced in individual flasks grown under standard conditions. Error bars represent the 95% confidence interval. The sample size are 5 flasks for the BUD_40_, and 6 flasks for the BUD_60_ and BUD_80_. **B)** Distribution of material area across all analyzed flasks. The sample size Source data are provided as a Source Data file.

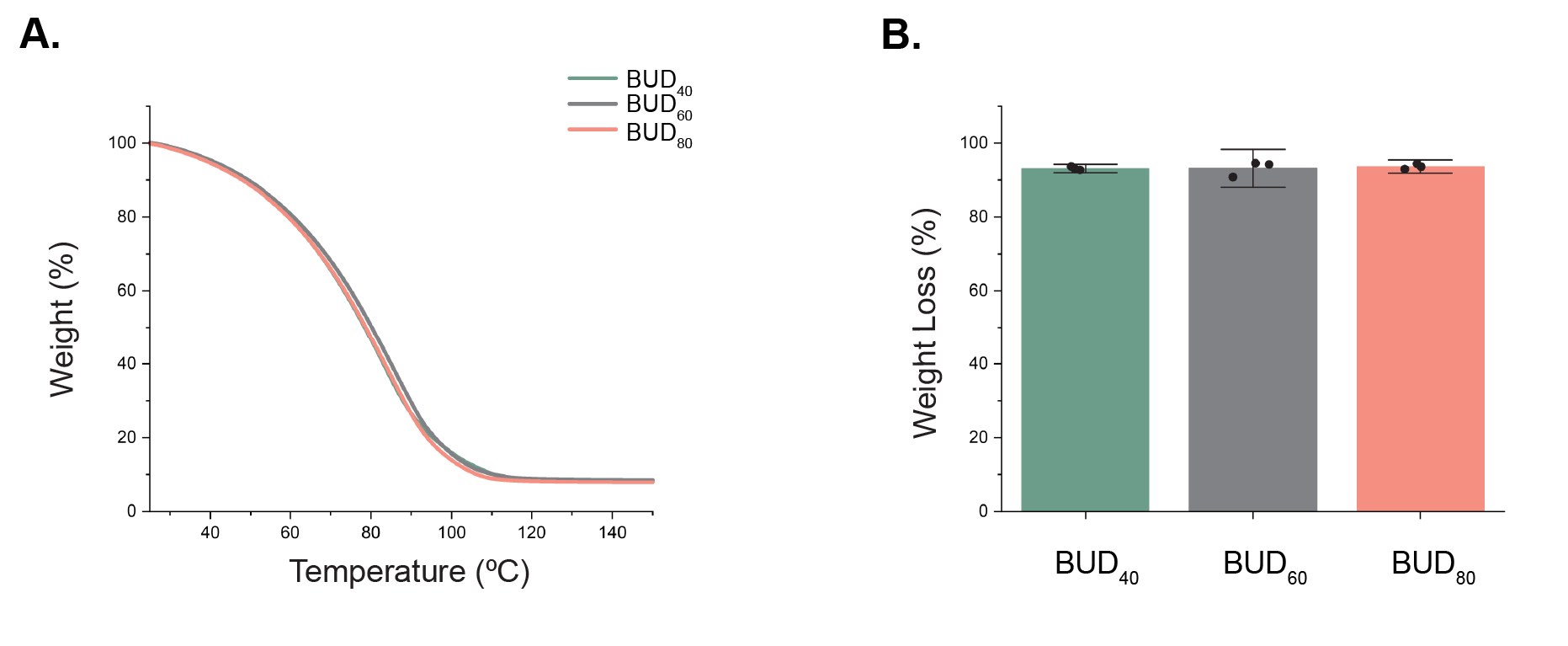
**Figure S2. The variant BUD-ELMs absorb similar amounts of water**. **A)** Thermogravimetric analysis experiments for each variant show weight % over a temperature range of 25°C-150°C. **B)** Total weight loss for the variant BUD-ELMs. Error bars are centered on the mean value and represent 95% confidence intervals for 3 samples. Source data are provided as a Source Data file.

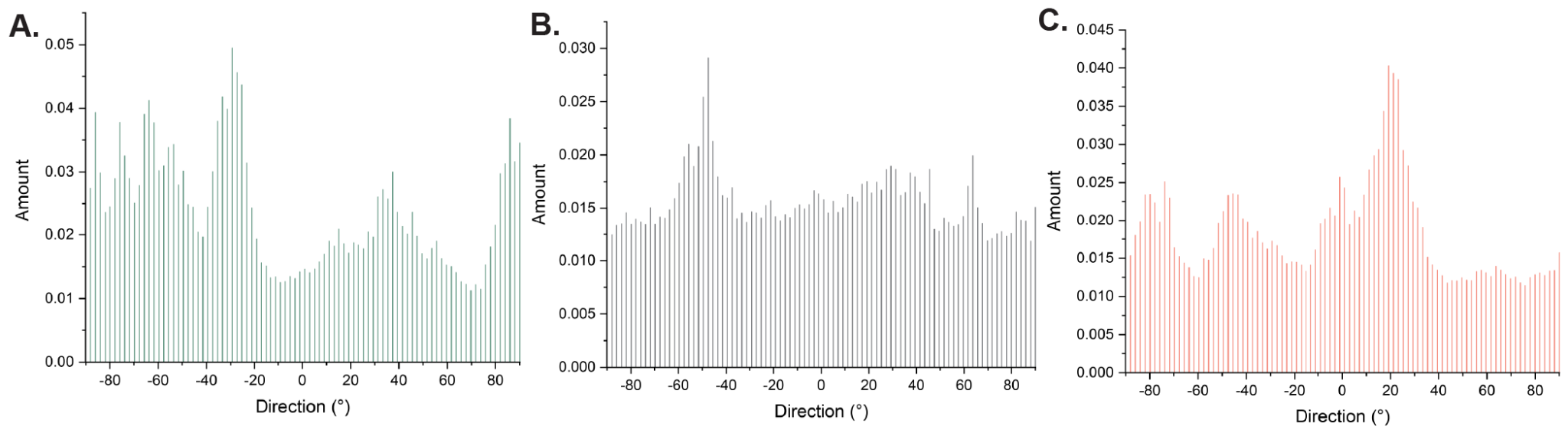
**Figure S3. The BUD_60_ variant fibers are more homogeneously aligned in all directionalities than the BUD_40_ and BUD_80_**. The amount of fibers aligned between -80° to 80° from the **A)** BUD_40_ microscopy images, **B)** BUD_60_ microscopy images, and **C)** BUD_80_ microscopy images. The sample size is 17 fibers.

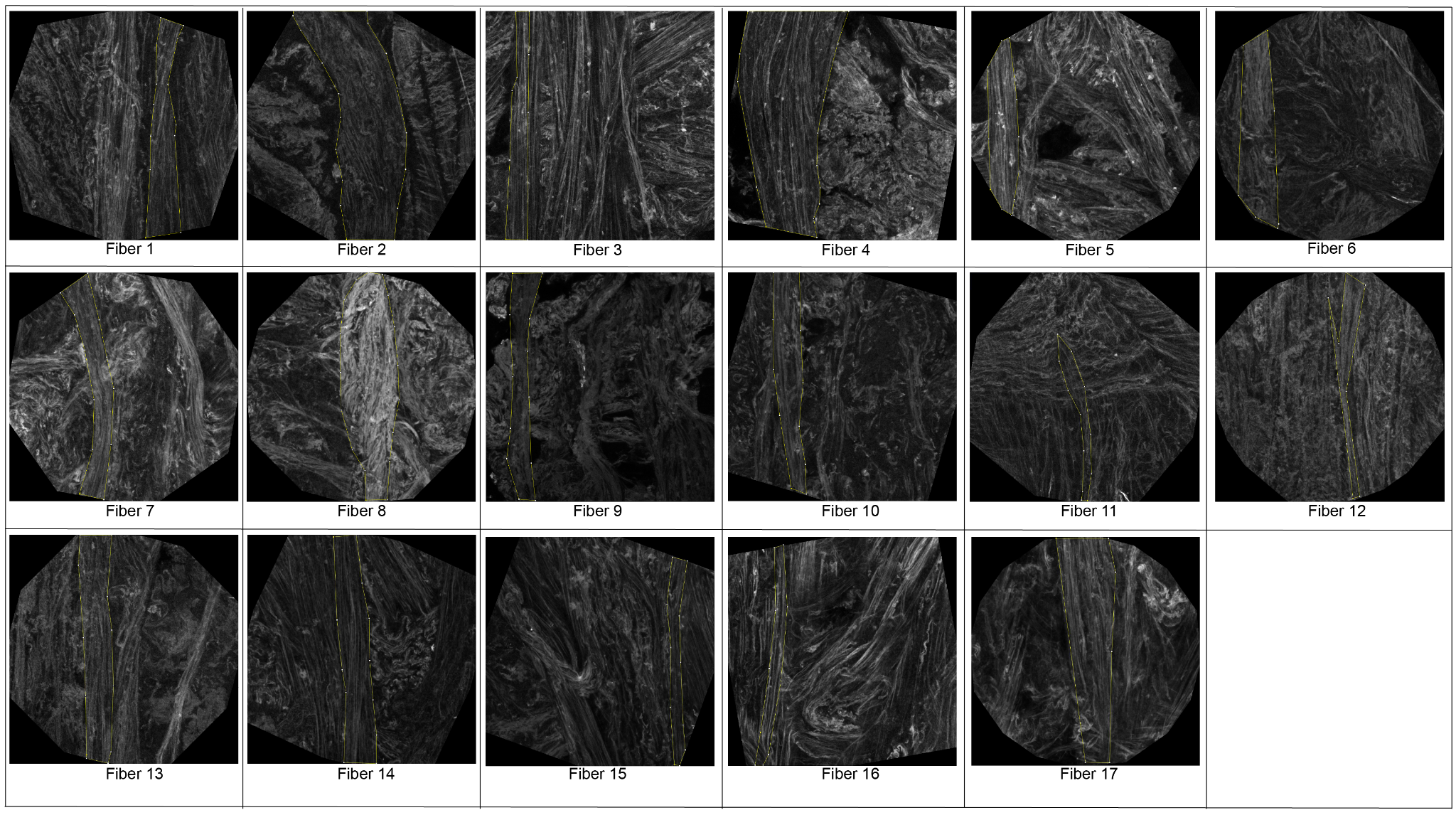

**Figure S4. Images of fibers analyzed for the BUD_40_.** The fibers that were measured are outlined in yellow. The sample size is 17 fibers.

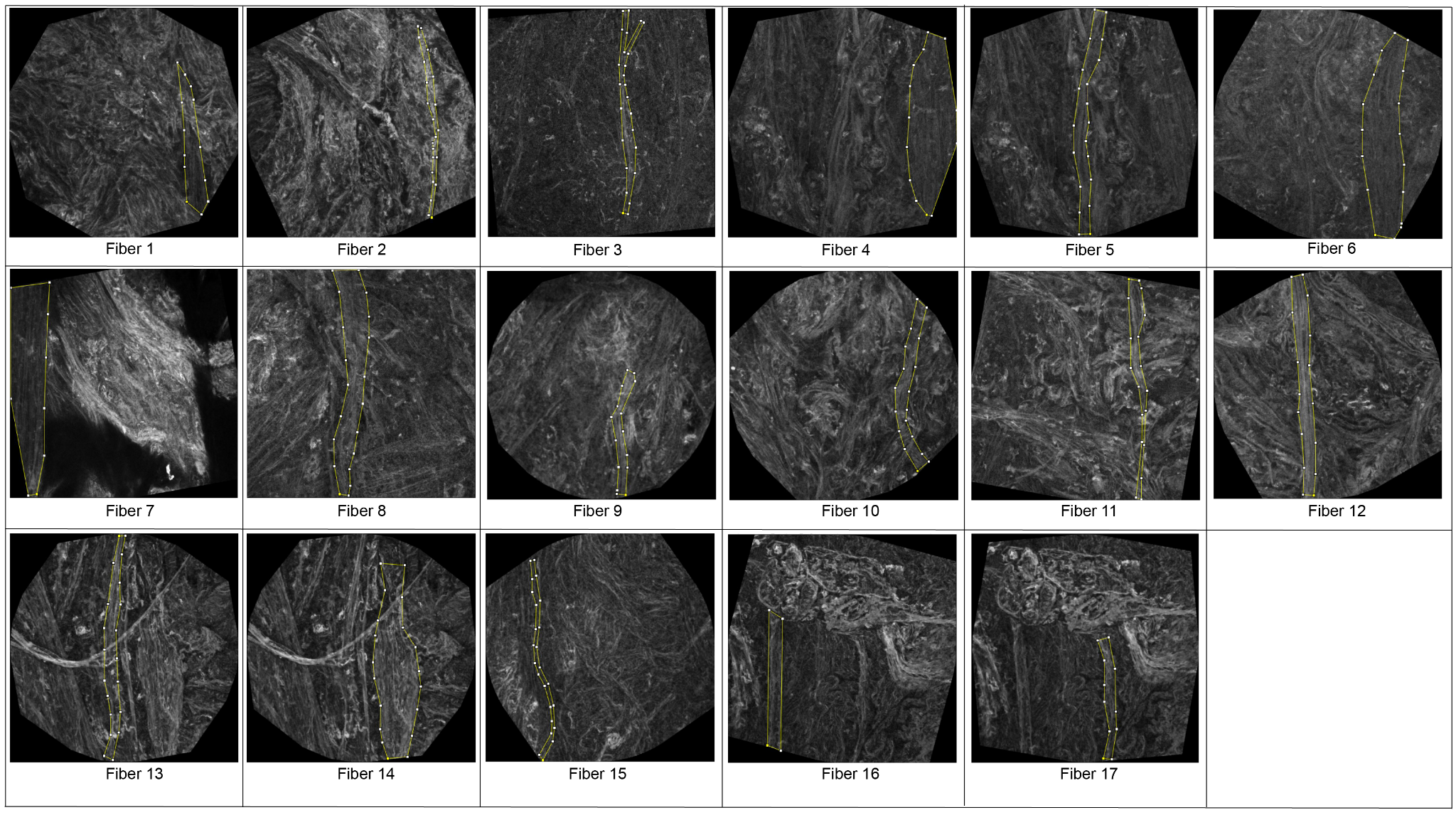

**Figure S5. Images of fibers analyzed for the BUD_60_.** The fibers that were measured are outlined in yellow. The sample size is 17 fibers.

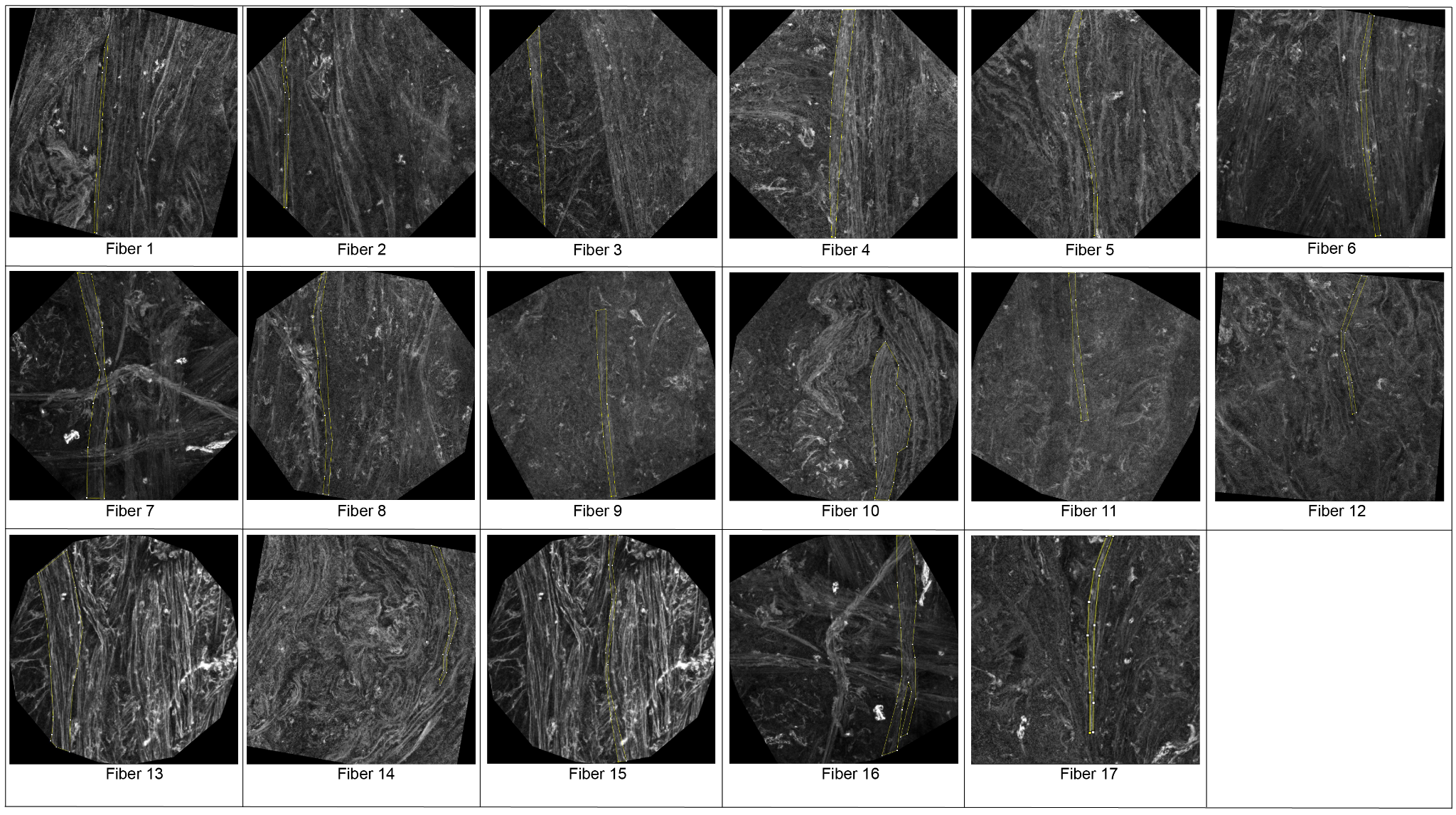

**Figure S6. Images of the fibers analyzed for the BUD_80_.** The fibers that were measured are outlined in yellow. The sample size is 17 fibers.

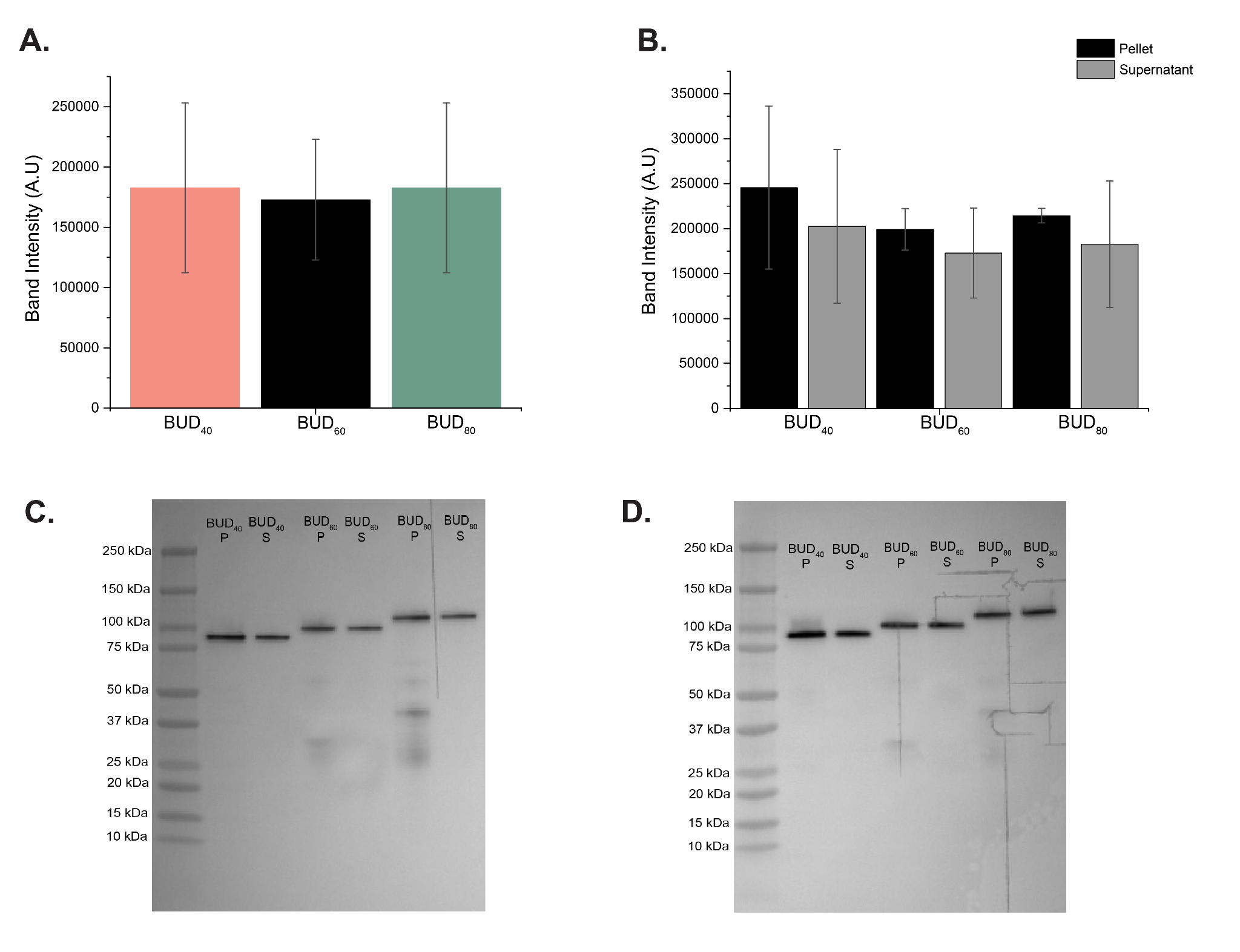
**Figure S7.** **The variants secrete similar amounts of protein**. The BUD-ELM variants were grown in standard conditions, harvested at 21 hrs, and immunoblot was performed using an anti-FLAG tag. **A)** The average band intensity for the supernatant of the BUD variants is plotted, showing similar protein concentrations. **B)** The pellet and supernatant of the BUD variants are shown, indicating no difference in the amount of protein on the surface of the cell and the amount being secreted. The sample size is 2. **C-D)** Immunoblot 1 and 2 showing protein secretion yields for the BUD variants**.** Lane 1 is the precision plus dual color standards ladder (BioRad), lane 2 is the BUD_40_ pellet fraction, lane 3 is the BUD_40_ supernatant fraction, lane 4 is the BUD_60_ pellet fraction, lane 5 is the BUD_60_ supernatant fractions, lane 6 is the BUD_80_ pellet fraction and lane 7 is the BUD_80_ supernatant fraction.

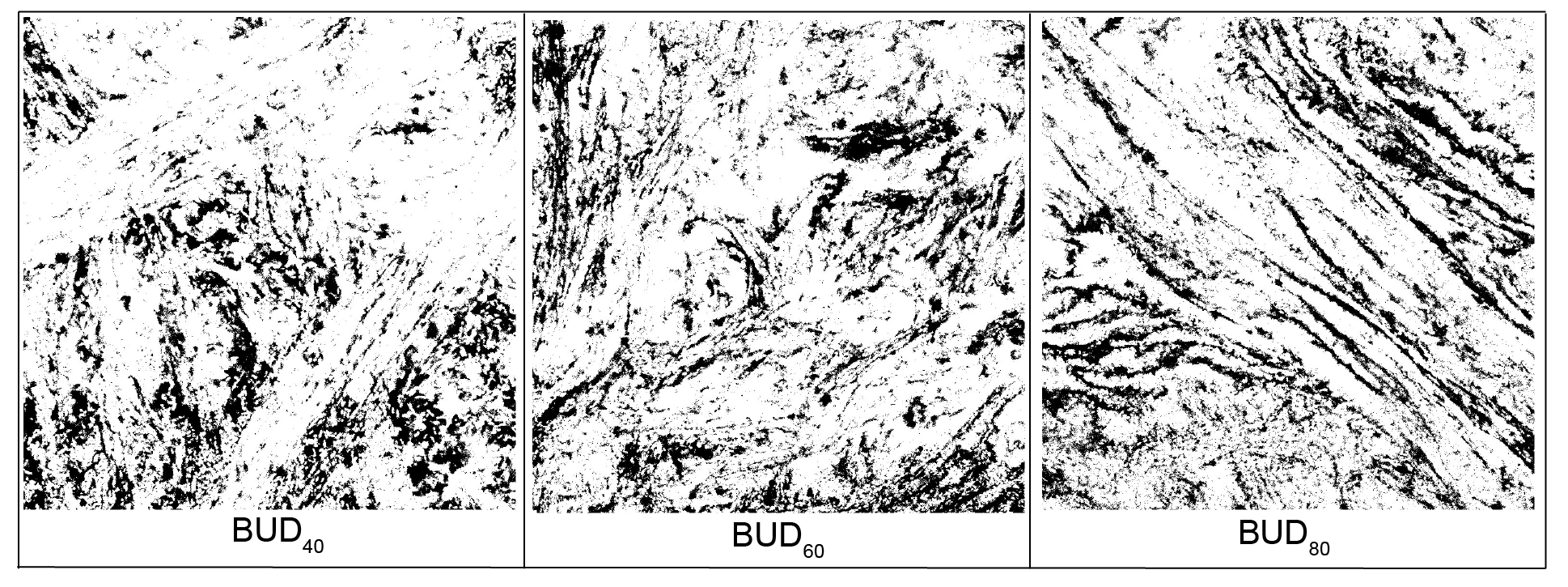
**Figure S8. Binary mask of the variant BUD-ELMs.** In white are the cell-rich areas, and in black are the non-cell-rich areas. The sample size is 27. Source data are provided as a Source Data file.

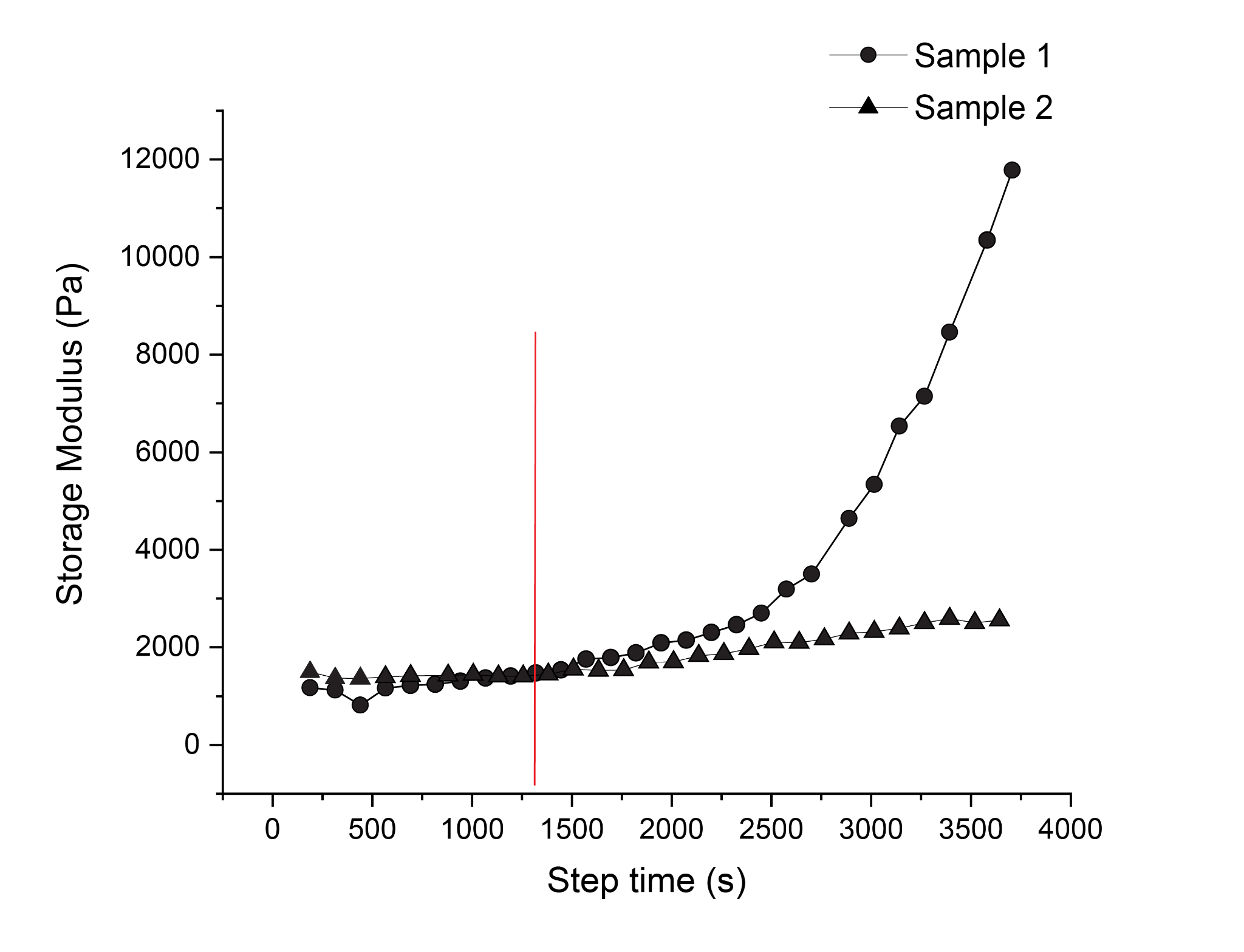

**Figure S9.** **Standard amplitude time sweeps for BUD_60_ show the materials start to dehydrate after 20 minutes as evidenced by the increase in storage modulus**. Standard oscillatory time sweeps were acquired for 3600s at a 0.5% oscillation strain and a fixed frequency of 0.1 rad/s for the BUD_60_ material. The vertical red line shows where the storage modulus starts to increase. Source data are provided as a Source Data file.

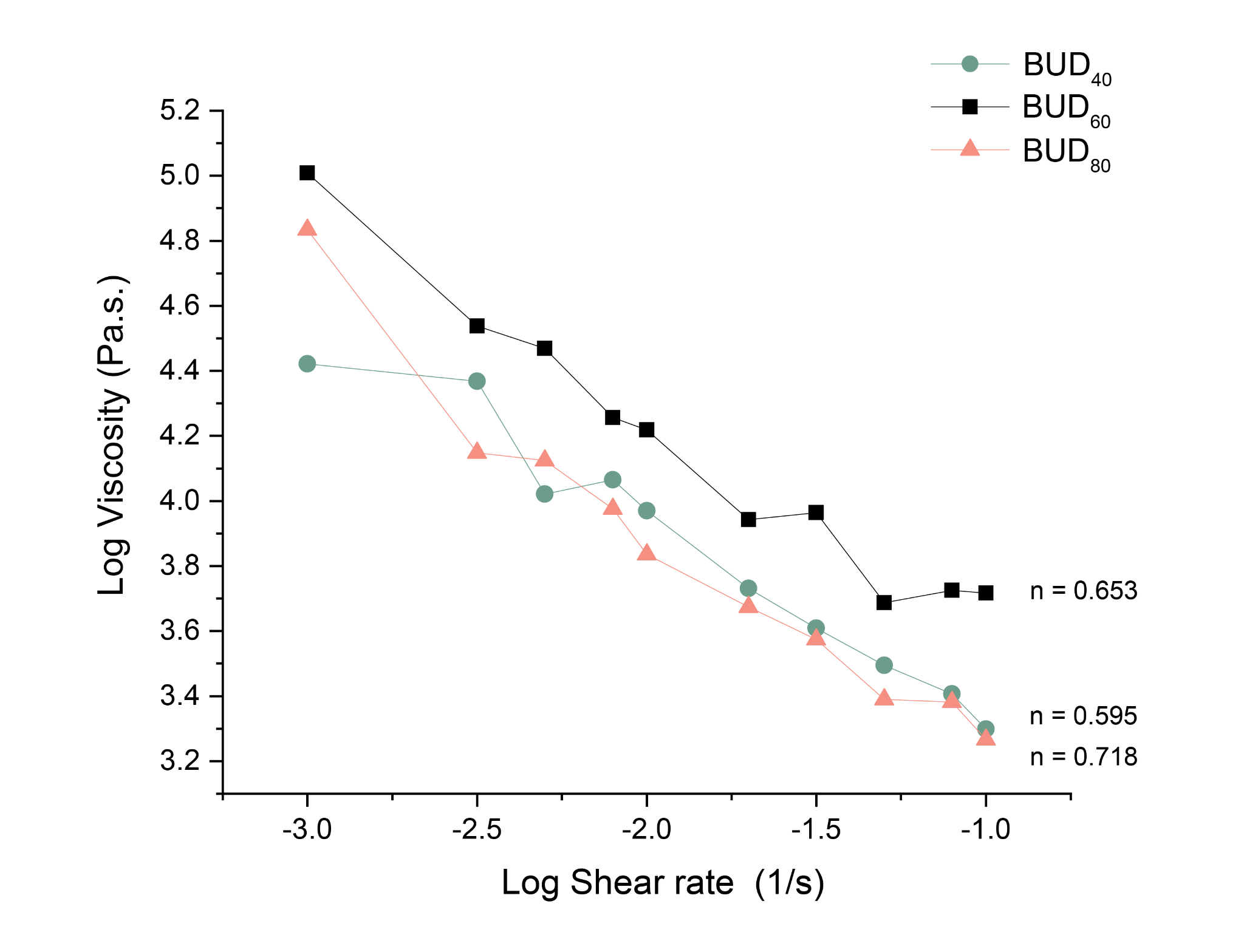

**Figure S10.** **The power law indexes for BUD variants are similar**. The materials were fitted to the power law model to obtain the power law index (n). This variable indicates the amount of shear-thinning behavior displayed by the BUD variants. The power law index for the BUD_40_, BUD_60_, and BUD_80_ is 0.653, 0.595, and 0.718 respectively, showing changing ELP length does not affect shear thinning behavior. Source data are provided as a Source Data file.

**Primers used to amplify the modified RsaA gene in the variant BUD-ELM strains**

**SMCAF087:** GATGCATACAAGCCCACGAAAG

**SMCAF092:** CTGAAGACCATGGCGGTGATCA

**Genscript ELP_40_ target sequence**

CGTCTCAAGTCGGTCTCATAGCGGAGGAGGCTCGGGTGACTACAAGGACGACGATGACAAGGTCCCAGGAGTTGGAGTCCCAGGAGGGGGCGTTCCGGGCGCAGGAGTTCCTGGAGTAGGAGTTCCAGGAGTGGGCGTGCCAGGGGTGGGCGTCCCAGGTGGGGGAGTTCCCGGAGCAGGTGTGCCTGGGGGCGGCGTGCCTGGAGTCGGAGTTCCGGGGGTGGGTGTACCGGGTGGAGGCGTACCAGGCGCGGGAGTGCCGGGCGTGGGCGTGCCAGGCGTCGGTGTACCCGGCGTTGGTGTTCCGGGCGGAGGTGTCCCCGGAGCTGGGGTTCCCGGTGGGGGTGTACCGGGCGTCGGGGTTCCCGGTGTGGGTGTCCCAGGTGGCGGCGTTCCCGGGGCGGGCGTACCTGGAGTGGGTGTGCCAGGAGTCGGCGTCCCAGGAGTCGGCGTACCAGGAGGTGGTGTTCCCGGGGCCGGAGTTCCCGGCGGAGGAGTTCCCGGCGTCGGCGTCCCTGGGGTCGGCGTCCCGGGAGGTGGAGTACCCGGAGCAGGAGTGCCGGGAGTCGGTGTACCTGGTGTCGGTGTCCCTGGTGTAGGTGTCCCGGGTGGTGGGGTGCCAGGTGCTGGCGTACCTGGGGGGGGGGTTCCTGGCGTAGGCGGCGGCTCGAGAGACCCAGTAGAGACG

**Genscript ELP_80_ target sequence**

CGTCTCAAGTCGGTCTCATAGCGGAGGAGGCTCGGGTGACTACAAGGACGACGATGACAAGGTCCCAGGAGTTGGAGTCCCAGGAGGGGGCGTTCCGGGCGCAGGAGTTCCTGGAGTAGGAGTTCCAGGAGTGGGCGTGCCAGGGGTGGGCGTCCCAGGTGGGGGAGTTCCCGGAGCAGGTGTGCCTGGGGGCGGCGTGCCTGGAGTCGGAGTTCCGGGGGTGGGTGTACCGGGTGGAGGCGTACCAGGCGCGGGAGTGCCGGGCGTGGGCGTGCCAGGCGTCGGTGTACCCGGCGTTGGTGTTCCGGGCGGAGGTGTCCCCGGAGCTGGGGTTCCCGGTGGGGGTGTACCGGGCGTCGGGGTTCCCGGTGTGGGTGTCCCAGGTGGCGGCGTTCCCGGGGCGGGCGTACCTGGAGTGGGTGTGCCAGGAGTCGGCGTCCCAGGAGTCGGCGTACCAGGAGGTGGTGTTCCCGGGGCCGGAGTTCCCGGCGGAGGAGTTCCCGGCGTCGGCGTCCCTGGGGTCGGCGTCCCGGGAGGTGGAGTACCCGGAGCAGGAGTGCCGGGAGTCGGTGTACCTGGTGTCGGTGTCCCTGGTGTAGGTGTCCCGGGTGGTGGGGTGCCAGGTGCTGGCGTACCTGGGGGGGGGGTTCCTGGCGTAGGCGTTCCGGGGGTGGGCGTTCCGGGCGGCGGGGTGCCGGGAGCAGGTGTCCCCGGCGTTGGTGTACCGGGGGTTGGTGTCCCAGGCGTAGGTGTGCCCGGTGGAGGGGTGCCGGGAGCTGGAGTGCCTGGAGGGGGTGTACCAGGGGTCGGTGTTCCCGGTGTAGGAGTACCGGGGGGCGGAGTCCCAGGAGCCGGCGTGCCGGGTGTTGGAGTCCCGGGAGTCGGAGTCCCTGGGGTAGGCGTTCCAGGGGGAGGGGTCCCCGGTGCAGGGGTTCCTGGCGGTGGTGTCCCAGGAGTTGGCGTCCCAGGAGTTGGAGTCCCAGGAGGGGGCGTTCCGGGCGCAGGAGTTCCTGGAGTAGGAGTTCCAGGAGTGGGCGTGCCAGGGGTGGGCGTCCCAGGTGGGGGAGTTCCCGGAGCAGGTGTGCCTGGGGGCGGCGTGCCTGGAGTCGGAGTTCCGGGGGTGGGTGTACCGGGTGGAGGCGTACCAGGCGCGGGAGTGCCGGGCGTGGGCGTGCCAGGCGTCGGTGTACCCGGCGTTGGTGTTCCGGGCGGAGGTGTCCCCGGAGCTGGGGTTCCCGGTGGGGGTGTACCGGGCGTCGGGGGCGGCTCGAGAGACCCAGTAGAGACG

**Genscript SpyTag target sequence**

GATTGATCTTACCCTTTAAGCGCGGGTATTGAACCAGGCTTATGCCCAGCATCGTTGCAAAGCTCATCGTCTCAAGTCGGTCTCACTCGGGAGGAGGCTCGGGTGCGCATATCGTAATGGTCGATGCATACAAGCCCACGAAAGGAGGTTCAGGCGGCGGAAGCGGTGGTGGAAGCGGAGGTGGGTCAGGCGGAGGCTCAGGGGGAGGTTCGGGTGGCGGGTCTAGAGACCCAGTAGAGACGGTAGCTAGCAGACTCAAACTAGATATATTATGCCCGCCATACAGACGAAACTAGTCGGA

**Table S1. List of the *C. crescentus* strains used in this work.**

| **Strain Name** | **Strain Designation** | **Source** | **genotype** |
| --- | --- | --- | --- |
| Original BUD-ELM | RCC002 | S. Molinari et al | NA1000 Δ*sapA::Pxyl mKate2,* *ΔrsaA(251-689)::* *ELP_60_-SpyTag* |
| BUD_40_ | gCAF018 | This work | NA1000 Δ*sapA::Pxyl mKate2,* *ΔrsaA(251-689)::* *ELP_40_-SpyTag* |
| BUD_80_ | gCAF019 | This work | NA1000 Δ*sapA::Pxyl mKate2,* *ΔrsaA(251-689)::* *ELP_80_-SpyTag* |

**Table S2. List of plasmids used in this work.**

| **Plasmid** | **Source** | **Description** | **Sequence** |
| --- | --- | --- | --- |
| pBTCAF008 | This work | Integration plasmid for strain building. *rsaA-N-terminus (800 bp), Golden Gate multicoloning site, rsaA-C-terminus (800 bp)* | GTTATCGGCATTTTCTTTTGCGTTTTTATTTGTTAACTGTTAATTGTCCTTGTTCAAGGATGCTGTCTTTGACAACAGATGTTTTCTTGCCTTTGATGTTCAGCAGGAAGCTCGGCGCAAACGTTGATTGTTTGTCTGCGTAGAATCCTCTGTTTGTCATATAGCTTGTAATCACGACATTGTTTCCTTTCGCTTGAGGTACAGCGAAGTGTGAGTAAGTAAAGGTTACATCGTTAGGATCAAGATCCATTTTTAACACAAGGCCAGTTTTGTTCAGCGGCTTGTATGGGCCAGTTAAAGAATTAGAAACATAACCAAGCATGTAAATATCGTTAGACGTAATGCCGTCAATCGTCATTTTTGATCCGCGGGAGTCAGTGAACAGATACCATTTGCCGTTCATTTTAAAGACGTTCGCGCGTTCAATTTCATCTGTTACTGTGTTAGATGCAATCAGCGGTTTCATCACTTTTTTCAGTGTGTAATCATCGTTTAGCTCAATCATACCGAGAGCGCCGTTTGCTAACTCAGCCGTGCGTTTTTTATCGCTTTGCAGAAGTTTTTGACTTTCTTGACGGAAGAATGATGTGCTTTTGCCATAGTATGCTTTGTTAAATAAAGATTCTTCGCCTTGGTAGCCATCTTCAGTTCCAGTGTTTGCTTCAAATACTAAGTATTTGTGGCCTTTATCTTCTACGTAGTGAGGATCTCTCAGCGTATGGTTGTCGCCTGAGCTGTAGTTGCCTTCATCGATGAACTGCTGTACATTTTGATACGTTTTTCCGTCACCGTCAAAGATTGATTTATAATCCTCTACACCGTTGATGTTCAAAGAGCTGTCTGATGCTGATACGTTAACTTGTGCAGTTGTCAGTGTTTGTTTGCCGTAATGTTTACCGGAGAAATCAGTGTAGAATAAACGGATTTTTCCGTCAGATGTAAATGTGGCTGAACCTGACCATTCTTGTGTTTGGTCTTTTAGGATAGAATCATTTGCATCGAATTTGTCGCTGTCTTTAAAGACGCGGCCAGCGTTTTTCCAGCTGTCAATAGAAGTTTCGCCGACTTTTTGATAGAACATGTAAATCGATGTGTCATCCGCATTTTTAGGATCTCCGGCTAATGCAAAGACGATGTGGTAGCCGTGATAGTTTGCGACAGTGCCGTCAGCGTTTTGTAATGGCCAGCTGTCCCAAACGTCCAGGCCTTTTGCAGAAGAGATATTTTTAATTGTGGACGAATCGAACTCAGGAACTTGATATTTTTCATTTTTTTGCTGTTCAGGGATTTGCAGCATATCATGGCGTGTAATATGGGAAATGCCGTATGTTTCCTTATATGGCTTTTGGTTCGTTTCTTTCGCAAACGCTTGAGTTGCGCCTCCTGCCAGCAGTGCGGTAGTAAAGGTTAATACTGTTGCTTGTTTTGCAAACTTTTTGATGTTCATCGTTCATGTCTCCTTTTTTATGTACTGTGTTAGCGGTCTGCTTCTTCCAGCCCTCCTGTTTGAAGATGGCAAGTTAGTTACGCACAATAAAAAAAGACCTAAAATATGTAAGGGGTGACGCCAAAGTATACACTTTGCCCTTTACACATTTTAGGTCTTGCCTGCTTTATCAGTAACAAACCCGCGCGATTTACTTTTCGACCTCATTCTATTAGACTCTCGTTTGGATTGCAACTGGTCTATTTTCCTCTTTTGTTTGATAGAAAATCATAAAAGGATTTGCAGACTACGGGCCTAAAGAACTAAAAAATCTATCTGTTTCTTTTCATTCTCTGTATTTTTTATAGTTTCTGTTGCATGGGCATAAAGTTGCCTTTTTAATCACAATTCAGAAAATATCATAATATCTCATTTCACTAAATAATAGAGCTTGGCCGGCGCCGTCCCGTCAAGTCAGCGTAATGCTCTGCCAGTGTTACAACCAATTAACCAATTCTGATTAGAAAAACTCATCGAGCATCAAATGAAACTGCAATTTATTCATATCAGGATTATCAATACCATATTTTTGAAAAAGCCGTTTCTGTAATGAAGGAGAAAACTCACCGAGGCAGTTCCATAGGATGGCAAGATCCTGGTATCGGTCTGCGATTCCGACTCGTCCAACATCAATACAACCTATTAATTTCCCCTCGTCAAAAATAAGGTTATCAAGTGAGAAATCACCATGAGTGACGACTGAATCCGGTGAGAATGGCAAAAACTTATGCATTTCTTTCCAGACTTGTTCAACAGGCCAACCATTACGCTCGTCATCAAAATCACTCGCATCAACCAAACCGTTATTCATTCGTGATTGCGCCTGAGCGAGACGAAATACGCGATCGCTGTTAAAAGGACAATTACAAACAGGAATCGAATGCAACCGGCGCAGGAACACTGCCAGCGCATCAACAATATTTTCACCTGAATCAGGATATTCTTCTAATACCTGGAATGCTGTTTTACCGGGGATCGCAGTGGTGAGTAACCATGCATCATCAGGAGTACGGATAAAATGCTTGATGGTCGGAAGAGGCATAAATTCCGTCAGCCAGTTTAGTCTGACCATCTCATCTGTAACATCATTGGCAACGCTACCTTTGCCATGTTTCAGAAACAACTCTGGCGCATCGGGCTTCCCATACAATCGATAGATTGTCGCACCTGATTGCCCGACATTATCGCGAGCCCATTTATACCCATATAAATCAGCATCCATGTTGGAATTTAATCGCGGCCTCGAGCAAGACGTTTCCCGTTGAATATGGCTCATAACACCCCTTGTATTACTGTTTATGTAAGCAGACAGTTTTATTGTTCATGATGATATATTTTTATCTTGTGCAATGTAACCCGGCCAAGCTCTAGAAATTGTAAACGTTAATATTTTGTTAAAATTCGCGTTAAATTTTTGTTAAATCAGCTCATTTTTTAACCAATAGGCCGAAATCGGCAAAATCCCTTATAAATCAAAAGAATAGCCCGAGATAGGGTTGAGTGTTGTTCCAGTTTGGAACAAGAGTCCACTATTAAAGAACGTGGACTCCAACGTCAAAGGGCGAAAAACCGTCTATCAGGGCGATGGCCCACTACGTGAACCATCACCCAAATCAAGTTTTTTGGGGTCGAGGTGCCGTAAAGCACTAAATCGGAACCCTAAAGGGAGCCCCCGATTTAGAGCTTGACGGGGAAAGCGAACGTGGCGAGAAAGGAAGGGAAGAAAGCGAAAGGAGCGGGCGCTAGGGCGCTGGCAAGTGTAGCGGTCACGCTGCGCGTAACCACCACACCCGCCGCGCTTAATGCGCCGCTACAGGGCGCGTAAAAGGATCTAGGTGAAGATCCTTTTTGATAATCTCATGGGGGATCGGTCTTGCCTTGCTCGTCGGTGATGTACTTCACCAGCTCCGCGAAGTCGCTCTTCTTGATGGAGCGCATGGGGACGTGCTTGGCAATCACGCGCACCCCCCGGCCGTTTTAGCGGCTAAAAAAGTCATGGCTCTGCCCTCGGGCGGACCACGCCCATCATGACCTTGCCAAGCTCGTCCTGCTTCTCTTCGATCTTCGCCAGCAGGGCGAGGATCGTGGCATCACCGAACCGCGCCGTGCGCGGGTCGTCGGTGAGCCAGAGTTTCAGCAGGCCGCCCAGGCGGCCCAGGTCGCCATTGATGCGGGCCAGCTCGCGGACGTGCTCATAGTCCACGACGCCCGTGATTTTGTAGCCCTGGCCGACGGCCAGCAGGTAGGCCGACAGGCTCATGCCGGCCGCCGCCGCCTTTTCCTCAATCGCTCTTCGTTCGTCTGGAAGGCAGTACACCTTGATAGGTGGGCTGCCCTTCCTGGTTGGCTTGGTTTCATCAGCCATCCGCTTGCCCTCATCTGTTACGCCGGCGGTAGCCGGCCAGCCTCGCAGAGCAGGATTCCCGTTGAGCACCGCCAGGTGCGAATAAGGGACAGTGAAGAAGGAACACCCGCTCGCGGGTGGGCCTACTTCACCTATCCTGCCCGGCTGACGCCGTTGGATACACCAAGGAAAGTCTACACGAACCCTTTGGCAAAATCCTGTATATCGTGCGAAAAAGGATGGATATACCGAAAAAATCGCTATAATGACCCCGAAGCAGGGTTATGCAGCGGAAAAGATCCGTCCATGACCAAAATCCCTTAACGTGAGTTTTCGTTCCACTGAGCGTCAGACCCCGTAGAAAAGATCAAAGGATCTTCTTGAGATCCTTTTTTTCTGCGCGTAATCTGCTGCTTGCAAACAAAAAAACCACCGCTACCAGCGGTGGTTTGTTTGCCGGATCAAGAGCTACCAACTCTTTTTCCGAAGGTAACTGGCTTCAGCAGAGCGCAGATACCAAATACTGTTCTTCTAGTGTAGCCGTAGTTAGGCCACCACTTCAAGAACTCTGTAGCACCGCCTACATACCTCGCTCTGCTAATCCTGTTACCAGTGGCTGCTGCCAGTGGCGATAAGTCGTGTCTTACCGGGTTGGACTCAAGACGATAGTTACCGGATAAGGCGCAGCGGTCGGGCTGAACGGGGGGTTCGTGCACACAGCCCAGCTTGGAGCGAACGACCTACACCGAACTGAGATACCTACAGCGTGAGCTATGAGAAAGCGCCACGCTTCCCGAAGGGAGAAAGGCGGACAGGTATCCGGTAAGCGGCAGGGTCGGAACAGGAAGAGCGCACGAGGGAGCTTCCAGGGGGAAACGCCTGGTATCTTTATAGTCCTGTCGGGTTTCGCCACCTCTGACTTGAGCGTCGATTTTTGTGATGCTCGTCAGGGGGGCGGAGCCTATGGAAAAACGCCAGCAACGCGGCCTTTTTACGGTTCCTGGCCTTTTGCTGGCCTTTTGCTCACATGTAATGTGAGTTAGCTCACTCATTAGGCACCCCAGGCTTTACACTTTATGCTTCCGGCTCGTATGTTGTGTGGAATTGTGAGCGGATAACAATTTCACACAGGAAACAGCTATGACCATGATTACGCCAAGCTACGTAATACGACTCACTAGTGGGGCCCgcgccactcggtcgcagggggtgtgggattttttttgggagacaatcctcatggcctatacgacggcccagttggtgactgcgtacaccaacgccaacctcggcaaggcgcctgacgccgccaccacgctgacgctcgacgcgtacgcgactcaaacccagacgggcggcctctcggacgccgctgcgctgaccaacaccctgaagctggtcaacagcacgacggctgttgccatccagacctaccagttcttcaccggcgttgccccgtcggccgctggtctggacttcctggtcgactcgaccaccaacaccaacgacctgaacgacgcgtactactcgaagttcgctcaggaaaaccgcttcatcaacttctcgatcaacctggccacgggcgccggcgccggcgcgacggctttcgccgccgcctacacgggcgtttcgtacgcccagacggtcgccaccgcctatgacaagatcatcggcaacgccgtcgcgaccgccgctggcgtcgacgtcgcggccgccgtggctttcctgagccgccaggccaacatcgactacctgaccgccttcgtgcgcgccaacacgccgttcacggccgctgccgacatcgatctggccgtcaaggccgccctgatcggcaccatcctgaacgccgccacggtgtcgggcatcggtggttacgcgaccgccacggccgcgatgatcaacgacctgtcggacggcgccctgtcgaccgacaacgcggctggcgtgaacctgttcaccgcctatccgtcgtcgggcgtgtcgggttcgGGCGGTAGCAGAGACCACTGGGTCTCAGTCTGGAGGAGGCTCGGGTGCTGACCCGGCCTTCGGCGGCTTCGAAACCCTCCGCGTCGCTGGCGCGGCGGCTCAAGGCTCGCACAACGCCAACGGCTTCACGGCTCTGCAACTGGGCGCGACGGCGGGTGCGACGACCTTCACCAACGTTGCGGTGAATGTCGGCCTGACCGTTCTGGCGGCTCCGACCGGTACGACGACCGTGACCCTGGCCAACGCCACGGGCACCTCGGACGTGTTCAACCTGACCCTGTCGTCCTCGGCCGCTCTGGCCGCTGGTACGGTTGCGCTGGCTGGCGTCGAGACGGTGAACATCGCCGCCACCGACACCAACACGACCGCTCACGTCGACACGCTGACGCTGCAAGCCACCTCGGCCAAGTCGATCGTGGTGACGGGCAACGCCGGTCTGAACCTGACCAACACCGGCAACACGGCTGTCACCAGCTTCGACGCCAGCGCCGTCACCGGCACGGGCTCGGCTGTGACCTTCGTGTCGGCCAACACCACGGTGGGTGAAGTCGTCACGATCCGCGGCGGCGCTGGCGCCGACTCGCTGACCGGTTCGGCCACCGCCAATGACACCATCATCGGTGGCGCTGGCGCTGACACCCTGGTCTACACCGGCGGTACGGACACCTTCACGGGTGGCACGGGCGCGGATATCTTCGATATCAACGCTATCGGCACCTCGACCGCTTTCGTGACGATCACCGACGCCGCTGTCGGCGACAAGCTCGACCTCGTCGGCATCTCGACGAACGGCGCTATCGCTGACGGCGCCTTCGGCGCTGCGGTCACCCTGGGCGCTGCTGCGACGCTAGCTTCGGCCGTGACGCGTCTCCGGATGTACAGGCATGCGTCGACCCTCTAGTCAAGGCCTTAAGTGAGTCGTATTACGGACTGGCCGTCGTTTTACAACGTCGTGACTGGGAAAACCCTGGCGTTACCCAACTTAATCGCCTTGCAGCACATCCCCCTTTCGCCAGCTGGCGTAATAGCGAAGAGGCCCGCACCGATCGCCCTTCCCAACAGTTGCGCAGCCTGAATGGCGAATGGCGCTTCGCTTGGTAATAAAGCCCGCTTCGGCGGGCTTTTTTTT |
| pCAF216 | This work | Integration plasmid for strain gCAF018. *rsaA-N-terminus (800 bp), ELP_40_, SpyTag, rsaA-C-terminus (800 bp)* | GTTATCGGCATTTTCTTTTGCGTTTTTATTTGTTAACTGTTAATTGTCCTTGTTCAAGGATGCTGTCTTTGACAACAGATGTTTTCTTGCCTTTGATGTTCAGCAGGAAGCTCGGCGCAAACGTTGATTGTTTGTCTGCGTAGAATCCTCTGTTTGTCATATAGCTTGTAATCACGACATTGTTTCCTTTCGCTTGAGGTACAGCGAAGTGTGAGTAAGTAAAGGTTACATCGTTAGGATCAAGATCCATTTTTAACACAAGGCCAGTTTTGTTCAGCGGCTTGTATGGGCCAGTTAAAGAATTAGAAACATAACCAAGCATGTAAATATCGTTAGACGTAATGCCGTCAATCGTCATTTTTGATCCGCGGGAGTCAGTGAACAGATACCATTTGCCGTTCATTTTAAAGACGTTCGCGCGTTCAATTTCATCTGTTACTGTGTTAGATGCAATCAGCGGTTTCATCACTTTTTTCAGTGTGTAATCATCGTTTAGCTCAATCATACCGAGAGCGCCGTTTGCTAACTCAGCCGTGCGTTTTTTATCGCTTTGCAGAAGTTTTTGACTTTCTTGACGGAAGAATGATGTGCTTTTGCCATAGTATGCTTTGTTAAATAAAGATTCTTCGCCTTGGTAGCCATCTTCAGTTCCAGTGTTTGCTTCAAATACTAAGTATTTGTGGCCTTTATCTTCTACGTAGTGAGGATCTCTCAGCGTATGGTTGTCGCCTGAGCTGTAGTTGCCTTCATCGATGAACTGCTGTACATTTTGATACGTTTTTCCGTCACCGTCAAAGATTGATTTATAATCCTCTACACCGTTGATGTTCAAAGAGCTGTCTGATGCTGATACGTTAACTTGTGCAGTTGTCAGTGTTTGTTTGCCGTAATGTTTACCGGAGAAATCAGTGTAGAATAAACGGATTTTTCCGTCAGATGTAAATGTGGCTGAACCTGACCATTCTTGTGTTTGGTCTTTTAGGATAGAATCATTTGCATCGAATTTGTCGCTGTCTTTAAAGACGCGGCCAGCGTTTTTCCAGCTGTCAATAGAAGTTTCGCCGACTTTTTGATAGAACATGTAAATCGATGTGTCATCCGCATTTTTAGGATCTCCGGCTAATGCAAAGACGATGTGGTAGCCGTGATAGTTTGCGACAGTGCCGTCAGCGTTTTGTAATGGCCAGCTGTCCCAAACGTCCAGGCCTTTTGCAGAAGAGATATTTTTAATTGTGGACGAATCGAACTCAGGAACTTGATATTTTTCATTTTTTTGCTGTTCAGGGATTTGCAGCATATCATGGCGTGTAATATGGGAAATGCCGTATGTTTCCTTATATGGCTTTTGGTTCGTTTCTTTCGCAAACGCTTGAGTTGCGCCTCCTGCCAGCAGTGCGGTAGTAAAGGTTAATACTGTTGCTTGTTTTGCAAACTTTTTGATGTTCATCGTTCATGTCTCCTTTTTTATGTACTGTGTTAGCGGTCTGCTTCTTCCAGCCCTCCTGTTTGAAGATGGCAAGTTAGTTACGCACAATAAAAAAAGACCTAAAATATGTAAGGGGTGACGCCAAAGTATACACTTTGCCCTTTACACATTTTAGGTCTTGCCTGCTTTATCAGTAACAAACCCGCGCGATTTACTTTTCGACCTCATTCTATTAGACTCTCGTTTGGATTGCAACTGGTCTATTTTCCTCTTTTGTTTGATAGAAAATCATAAAAGGATTTGCAGACTACGGGCCTAAAGAACTAAAAAATCTATCTGTTTCTTTTCATTCTCTGTATTTTTTATAGTTTCTGTTGCATGGGCATAAAGTTGCCTTTTTAATCACAATTCAGAAAATATCATAATATCTCATTTCACTAAATAATAGAGCTTGGCCGGCGCCGTCCCGTCAAGTCAGCGTAATGCTCTGCCAGTGTTACAACCAATTAACCAATTCTGATTAGAAAAACTCATCGAGCATCAAATGAAACTGCAATTTATTCATATCAGGATTATCAATACCATATTTTTGAAAAAGCCGTTTCTGTAATGAAGGAGAAAACTCACCGAGGCAGTTCCATAGGATGGCAAGATCCTGGTATCGGTCTGCGATTCCGACTCGTCCAACATCAATACAACCTATTAATTTCCCCTCGTCAAAAATAAGGTTATCAAGTGAGAAATCACCATGAGTGACGACTGAATCCGGTGAGAATGGCAAAAACTTATGCATTTCTTTCCAGACTTGTTCAACAGGCCAACCATTACGCTCGTCATCAAAATCACTCGCATCAACCAAACCGTTATTCATTCGTGATTGCGCCTGAGCGAGACGAAATACGCGATCGCTGTTAAAAGGACAATTACAAACAGGAATCGAATGCAACCGGCGCAGGAACACTGCCAGCGCATCAACAATATTTTCACCTGAATCAGGATATTCTTCTAATACCTGGAATGCTGTTTTACCGGGGATCGCAGTGGTGAGTAACCATGCATCATCAGGAGTACGGATAAAATGCTTGATGGTCGGAAGAGGCATAAATTCCGTCAGCCAGTTTAGTCTGACCATCTCATCTGTAACATCATTGGCAACGCTACCTTTGCCATGTTTCAGAAACAACTCTGGCGCATCGGGCTTCCCATACAATCGATAGATTGTCGCACCTGATTGCCCGACATTATCGCGAGCCCATTTATACCCATATAAATCAGCATCCATGTTGGAATTTAATCGCGGCCTCGAGCAAGACGTTTCCCGTTGAATATGGCTCATAACACCCCTTGTATTACTGTTTATGTAAGCAGACAGTTTTATTGTTCATGATGATATATTTTTATCTTGTGCAATGTAACCCGGCCAAGCTCTAGAAATTGTAAACGTTAATATTTTGTTAAAATTCGCGTTAAATTTTTGTTAAATCAGCTCATTTTTTAACCAATAGGCCGAAATCGGCAAAATCCCTTATAAATCAAAAGAATAGCCCGAGATAGGGTTGAGTGTTGTTCCAGTTTGGAACAAGAGTCCACTATTAAAGAACGTGGACTCCAACGTCAAAGGGCGAAAAACCGTCTATCAGGGCGATGGCCCACTACGTGAACCATCACCCAAATCAAGTTTTTTGGGGTCGAGGTGCCGTAAAGCACTAAATCGGAACCCTAAAGGGAGCCCCCGATTTAGAGCTTGACGGGGAAAGCGAACGTGGCGAGAAAGGAAGGGAAGAAAGCGAAAGGAGCGGGCGCTAGGGCGCTGGCAAGTGTAGCGGTCACGCTGCGCGTAACCACCACACCCGCCGCGCTTAATGCGCCGCTACAGGGCGCGTAAAAGGATCTAGGTGAAGATCCTTTTTGATAATCTCATGGGGGATCGGTCTTGCCTTGCTCGTCGGTGATGTACTTCACCAGCTCCGCGAAGTCGCTCTTCTTGATGGAGCGCATGGGGACGTGCTTGGCAATCACGCGCACCCCCCGGCCGTTTTAGCGGCTAAAAAAGTCATGGCTCTGCCCTCGGGCGGACCACGCCCATCATGACCTTGCCAAGCTCGTCCTGCTTCTCTTCGATCTTCGCCAGCAGGGCGAGGATCGTGGCATCACCGAACCGCGCCGTGCGCGGGTCGTCGGTGAGCCAGAGTTTCAGCAGGCCGCCCAGGCGGCCCAGGTCGCCATTGATGCGGGCCAGCTCGCGGACGTGCTCATAGTCCACGACGCCCGTGATTTTGTAGCCCTGGCCGACGGCCAGCAGGTAGGCCGACAGGCTCATGCCGGCCGCCGCCGCCTTTTCCTCAATCGCTCTTCGTTCGTCTGGAAGGCAGTACACCTTGATAGGTGGGCTGCCCTTCCTGGTTGGCTTGGTTTCATCAGCCATCCGCTTGCCCTCATCTGTTACGCCGGCGGTAGCCGGCCAGCCTCGCAGAGCAGGATTCCCGTTGAGCACCGCCAGGTGCGAATAAGGGACAGTGAAGAAGGAACACCCGCTCGCGGGTGGGCCTACTTCACCTATCCTGCCCGGCTGACGCCGTTGGATACACCAAGGAAAGTCTACACGAACCCTTTGGCAAAATCCTGTATATCGTGCGAAAAAGGATGGATATACCGAAAAAATCGCTATAATGACCCCGAAGCAGGGTTATGCAGCGGAAAAGATCCGTCCATGACCAAAATCCCTTAACGTGAGTTTTCGTTCCACTGAGCGTCAGACCCCGTAGAAAAGATCAAAGGATCTTCTTGAGATCCTTTTTTTCTGCGCGTAATCTGCTGCTTGCAAACAAAAAAACCACCGCTACCAGCGGTGGTTTGTTTGCCGGATCAAGAGCTACCAACTCTTTTTCCGAAGGTAACTGGCTTCAGCAGAGCGCAGATACCAAATACTGTTCTTCTAGTGTAGCCGTAGTTAGGCCACCACTTCAAGAACTCTGTAGCACCGCCTACATACCTCGCTCTGCTAATCCTGTTACCAGTGGCTGCTGCCAGTGGCGATAAGTCGTGTCTTACCGGGTTGGACTCAAGACGATAGTTACCGGATAAGGCGCAGCGGTCGGGCTGAACGGGGGGTTCGTGCACACAGCCCAGCTTGGAGCGAACGACCTACACCGAACTGAGATACCTACAGCGTGAGCTATGAGAAAGCGCCACGCTTCCCGAAGGGAGAAAGGCGGACAGGTATCCGGTAAGCGGCAGGGTCGGAACAGGAAGAGCGCACGAGGGAGCTTCCAGGGGGAAACGCCTGGTATCTTTATAGTCCTGTCGGGTTTCGCCACCTCTGACTTGAGCGTCGATTTTTGTGATGCTCGTCAGGGGGGCGGAGCCTATGGAAAAACGCCAGCAACGCGGCCTTTTTACGGTTCCTGGCCTTTTGCTGGCCTTTTGCTCACATGTAATGTGAGTTAGCTCACTCATTAGGCACCCCAGGCTTTACACTTTATGCTTCCGGCTCGTATGTTGTGTGGAATTGTGAGCGGATAACAATTTCACACAGGAAACAGCTATGACCATGATTACGCCAAGCTACGTAATACGACTCACTAGTGGGGCCCgcgccactcggtcgcagggggtgtgggattttttttgggagacaatcctcatggcctatacgacggcccagttggtgactgcgtacaccaacgccaacctcggcaaggcgcctgacgccgccaccacgctgacgctcgacgcgtacgcgactcaaacccagacgggcggcctctcggacgccgctgcgctgaccaacaccctgaagctggtcaacagcacgacggctgttgccatccagacctaccagttcttcaccggcgttgccccgtcggccgctggtctggacttcctggtcgactcgaccaccaacaccaacgacctgaacgacgcgtactactcgaagttcgctcaggaaaaccgcttcatcaacttctcgatcaacctggccacgggcgccggcgccggcgcgacggctttcgccgccgcctacacgggcgtttcgtacgcccagacggtcgccaccgcctatgacaagatcatcggcaacgccgtcgcgaccgccgctggcgtcgacgtcgcggccgccgtggctttcctgagccgccaggccaacatcgactacctgaccgccttcgtgcgcgccaacacgccgttcacggccgctgccgacatcgatctggccgtcaaggccgccctgatcggcaccatcctgaacgccgccacggtgtcgggcatcggtggttacgcgaccgccacggccgcgatgatcaacgacctgtcggacggcgccctgtcgaccgacaacgcggctggcgtgaacctgttcaccgcctatccgtcgtcgggcgtgtcgggttcgGGCGGTAGCGGAGGAGGCTCGGGTGACTACAAGGACGACGATGACAAGGTCCCAGGAGTTGGAGTCCCAGGAGGGGGCGTTCCGGGCGCAGGAGTTCCTGGAGTAGGAGTTCCAGGAGTGGGCGTGCCAGGGGTGGGCGTCCCAGGTGGGGGAGTTCCCGGAGCAGGTGTGCCTGGGGGCGGCGTGCCTGGAGTCGGAGTTCCGGGGGTGGGTGTACCGGGTGGAGGCGTACCAGGCGCGGGAGTGCCGGGCGTGGGCGTGCCAGGCGTCGGTGTACCCGGCGTTGGTGTTCCGGGCGGAGGTGTCCCCGGAGCTGGGGTTCCCGGTGGGGGTGTACCGGGCGTCGGGGTTCCCGGTGTGGGTGTCCCAGGTGGCGGCGTTCCCGGGGCGGGCGTACCTGGAGTGGGTGTGCCAGGAGTCGGCGTCCCAGGAGTCGGCGTACCAGGAGGTGGTGTTCCCGGGGCCGGAGTTCCCGGCGGAGGAGTTCCCGGCGTCGGCGTCCCTGGGGTCGGCGTCCCGGGAGGTGGAGTACCCGGAGCAGGAGTGCCGGGAGTCGGTGTACCTGGTGTCGGTGTCCCTGGTGTAGGTGTCCCGGGTGGTGGGGTGCCAGGTGCTGGCGTACCTGGGGGGGGGGTTCCTGGCGTAGGCGGCGGCTCGGGAGGAGGCTCGGGTGCGCATATCGTAATGGTCGATGCATACAAGCCCACGAAAGGAGGTTCAGGCGGCGGAAGCGGTGGTGGAAGCGGAGGTGGGTCAGGCGGAGGCTCAGGGGGAGGTTCGGGTGGCGGGTCTGGAGGAGGCTCGGGTGCTGACCCGGCCTTCGGCGGCTTCGAAACCCTCCGCGTCGCTGGCGCGGCGGCTCAAGGCTCGCACAACGCCAACGGCTTCACGGCTCTGCAACTGGGCGCGACGGCGGGTGCGACGACCTTCACCAACGTTGCGGTGAATGTCGGCCTGACCGTTCTGGCGGCTCCGACCGGTACGACGACCGTGACCCTGGCCAACGCCACGGGCACCTCGGACGTGTTCAACCTGACCCTGTCGTCCTCGGCCGCTCTGGCCGCTGGTACGGTTGCGCTGGCTGGCGTCGAGACGGTGAACATCGCCGCCACCGACACCAACACGACCGCTCACGTCGACACGCTGACGCTGCAAGCCACCTCGGCCAAGTCGATCGTGGTGACGGGCAACGCCGGTCTGAACCTGACCAACACCGGCAACACGGCTGTCACCAGCTTCGACGCCAGCGCCGTCACCGGCACGGGCTCGGCTGTGACCTTCGTGTCGGCCAACACCACGGTGGGTGAAGTCGTCACGATCCGCGGCGGCGCTGGCGCCGACTCGCTGACCGGTTCGGCCACCGCCAATGACACCATCATCGGTGGCGCTGGCGCTGACACCCTGGTCTACACCGGCGGTACGGACACCTTCACGGGTGGCACGGGCGCGGATATCTTCGATATCAACGCTATCGGCACCTCGACCGCTTTCGTGACGATCACCGACGCCGCTGTCGGCGACAAGCTCGACCTCGTCGGCATCTCGACGAACGGCGCTATCGCTGACGGCGCCTTCGGCGCTGCGGTCACCCTGGGCGCTGCTGCGACGCTAGCTTCGGCCGTGACGCGTCTCCGGATGTACAGGCATGCGTCGACCCTCTAGTCAAGGCCTTAAGTGAGTCGTATTACGGACTGGCCGTCGTTTTACAACGTCGTGACTGGGAAAACCCTGGCGTTACCCAACTTAATCGCCTTGCAGCACATCCCCCTTTCGCCAGCTGGCGTAATAGCGAAGAGGCCCGCACCGATCGCCCTTCCCAACAGTTGCGCAGCCTGAATGGCGAATGGCGCTTCGCTTGGTAATAAAGCCCGCTTCGGCGGGCTTTTTTTT |
| pCAF215 | This work | Integration plasmid for strain gCAF019. *rsaA-N-terminus (800 bp), ELP_80_, SpyTag, rsaA-C-terminus (800 bp)* | GTTATCGGCATTTTCTTTTGCGTTTTTATTTGTTAACTGTTAATTGTCCTTGTTCAAGGATGCTGTCTTTGACAACAGATGTTTTCTTGCCTTTGATGTTCAGCAGGAAGCTCGGCGCAAACGTTGATTGTTTGTCTGCGTAGAATCCTCTGTTTGTCATATAGCTTGTAATCACGACATTGTTTCCTTTCGCTTGAGGTACAGCGAAGTGTGAGTAAGTAAAGGTTACATCGTTAGGATCAAGATCCATTTTTAACACAAGGCCAGTTTTGTTCAGCGGCTTGTATGGGCCAGTTAAAGAATTAGAAACATAACCAAGCATGTAAATATCGTTAGACGTAATGCCGTCAATCGTCATTTTTGATCCGCGGGAGTCAGTGAACAGATACCATTTGCCGTTCATTTTAAAGACGTTCGCGCGTTCAATTTCATCTGTTACTGTGTTAGATGCAATCAGCGGTTTCATCACTTTTTTCAGTGTGTAATCATCGTTTAGCTCAATCATACCGAGAGCGCCGTTTGCTAACTCAGCCGTGCGTTTTTTATCGCTTTGCAGAAGTTTTTGACTTTCTTGACGGAAGAATGATGTGCTTTTGCCATAGTATGCTTTGTTAAATAAAGATTCTTCGCCTTGGTAGCCATCTTCAGTTCCAGTGTTTGCTTCAAATACTAAGTATTTGTGGCCTTTATCTTCTACGTAGTGAGGATCTCTCAGCGTATGGTTGTCGCCTGAGCTGTAGTTGCCTTCATCGATGAACTGCTGTACATTTTGATACGTTTTTCCGTCACCGTCAAAGATTGATTTATAATCCTCTACACCGTTGATGTTCAAAGAGCTGTCTGATGCTGATACGTTAACTTGTGCAGTTGTCAGTGTTTGTTTGCCGTAATGTTTACCGGAGAAATCAGTGTAGAATAAACGGATTTTTCCGTCAGATGTAAATGTGGCTGAACCTGACCATTCTTGTGTTTGGTCTTTTAGGATAGAATCATTTGCATCGAATTTGTCGCTGTCTTTAAAGACGCGGCCAGCGTTTTTCCAGCTGTCAATAGAAGTTTCGCCGACTTTTTGATAGAACATGTAAATCGATGTGTCATCCGCATTTTTAGGATCTCCGGCTAATGCAAAGACGATGTGGTAGCCGTGATAGTTTGCGACAGTGCCGTCAGCGTTTTGTAATGGCCAGCTGTCCCAAACGTCCAGGCCTTTTGCAGAAGAGATATTTTTAATTGTGGACGAATCGAACTCAGGAACTTGATATTTTTCATTTTTTTGCTGTTCAGGGATTTGCAGCATATCATGGCGTGTAATATGGGAAATGCCGTATGTTTCCTTATATGGCTTTTGGTTCGTTTCTTTCGCAAACGCTTGAGTTGCGCCTCCTGCCAGCAGTGCGGTAGTAAAGGTTAATACTGTTGCTTGTTTTGCAAACTTTTTGATGTTCATCGTTCATGTCTCCTTTTTTATGTACTGTGTTAGCGGTCTGCTTCTTCCAGCCCTCCTGTTTGAAGATGGCAAGTTAGTTACGCACAATAAAAAAAGACCTAAAATATGTAAGGGGTGACGCCAAAGTATACACTTTGCCCTTTACACATTTTAGGTCTTGCCTGCTTTATCAGTAACAAACCCGCGCGATTTACTTTTCGACCTCATTCTATTAGACTCTCGTTTGGATTGCAACTGGTCTATTTTCCTCTTTTGTTTGATAGAAAATCATAAAAGGATTTGCAGACTACGGGCCTAAAGAACTAAAAAATCTATCTGTTTCTTTTCATTCTCTGTATTTTTTATAGTTTCTGTTGCATGGGCATAAAGTTGCCTTTTTAATCACAATTCAGAAAATATCATAATATCTCATTTCACTAAATAATAGAGCTTGGCCGGCGCCGTCCCGTCAAGTCAGCGTAATGCTCTGCCAGTGTTACAACCAATTAACCAATTCTGATTAGAAAAACTCATCGAGCATCAAATGAAACTGCAATTTATTCATATCAGGATTATCAATACCATATTTTTGAAAAAGCCGTTTCTGTAATGAAGGAGAAAACTCACCGAGGCAGTTCCATAGGATGGCAAGATCCTGGTATCGGTCTGCGATTCCGACTCGTCCAACATCAATACAACCTATTAATTTCCCCTCGTCAAAAATAAGGTTATCAAGTGAGAAATCACCATGAGTGACGACTGAATCCGGTGAGAATGGCAAAAACTTATGCATTTCTTTCCAGACTTGTTCAACAGGCCAACCATTACGCTCGTCATCAAAATCACTCGCATCAACCAAACCGTTATTCATTCGTGATTGCGCCTGAGCGAGACGAAATACGCGATCGCTGTTAAAAGGACAATTACAAACAGGAATCGAATGCAACCGGCGCAGGAACACTGCCAGCGCATCAACAATATTTTCACCTGAATCAGGATATTCTTCTAATACCTGGAATGCTGTTTTACCGGGGATCGCAGTGGTGAGTAACCATGCATCATCAGGAGTACGGATAAAATGCTTGATGGTCGGAAGAGGCATAAATTCCGTCAGCCAGTTTAGTCTGACCATCTCATCTGTAACATCATTGGCAACGCTACCTTTGCCATGTTTCAGAAACAACTCTGGCGCATCGGGCTTCCCATACAATCGATAGATTGTCGCACCTGATTGCCCGACATTATCGCGAGCCCATTTATACCCATATAAATCAGCATCCATGTTGGAATTTAATCGCGGCCTCGAGCAAGACGTTTCCCGTTGAATATGGCTCATAACACCCCTTGTATTACTGTTTATGTAAGCAGACAGTTTTATTGTTCATGATGATATATTTTTATCTTGTGCAATGTAACCCGGCCAAGCTCTAGAAATTGTAAACGTTAATATTTTGTTAAAATTCGCGTTAAATTTTTGTTAAATCAGCTCATTTTTTAACCAATAGGCCGAAATCGGCAAAATCCCTTATAAATCAAAAGAATAGCCCGAGATAGGGTTGAGTGTTGTTCCAGTTTGGAACAAGAGTCCACTATTAAAGAACGTGGACTCCAACGTCAAAGGGCGAAAAACCGTCTATCAGGGCGATGGCCCACTACGTGAACCATCACCCAAATCAAGTTTTTTGGGGTCGAGGTGCCGTAAAGCACTAAATCGGAACCCTAAAGGGAGCCCCCGATTTAGAGCTTGACGGGGAAAGCGAACGTGGCGAGAAAGGAAGGGAAGAAAGCGAAAGGAGCGGGCGCTAGGGCGCTGGCAAGTGTAGCGGTCACGCTGCGCGTAACCACCACACCCGCCGCGCTTAATGCGCCGCTACAGGGCGCGTAAAAGGATCTAGGTGAAGATCCTTTTTGATAATCTCATGGGGGATCGGTCTTGCCTTGCTCGTCGGTGATGTACTTCACCAGCTCCGCGAAGTCGCTCTTCTTGATGGAGCGCATGGGGACGTGCTTGGCAATCACGCGCACCCCCCGGCCGTTTTAGCGGCTAAAAAAGTCATGGCTCTGCCCTCGGGCGGACCACGCCCATCATGACCTTGCCAAGCTCGTCCTGCTTCTCTTCGATCTTCGCCAGCAGGGCGAGGATCGTGGCATCACCGAACCGCGCCGTGCGCGGGTCGTCGGTGAGCCAGAGTTTCAGCAGGCCGCCCAGGCGGCCCAGGTCGCCATTGATGCGGGCCAGCTCGCGGACGTGCTCATAGTCCACGACGCCCGTGATTTTGTAGCCCTGGCCGACGGCCAGCAGGTAGGCCGACAGGCTCATGCCGGCCGCCGCCGCCTTTTCCTCAATCGCTCTTCGTTCGTCTGGAAGGCAGTACACCTTGATAGGTGGGCTGCCCTTCCTGGTTGGCTTGGTTTCATCAGCCATCCGCTTGCCCTCATCTGTTACGCCGGCGGTAGCCGGCCAGCCTCGCAGAGCAGGATTCCCGTTGAGCACCGCCAGGTGCGAATAAGGGACAGTGAAGAAGGAACACCCGCTCGCGGGTGGGCCTACTTCACCTATCCTGCCCGGCTGACGCCGTTGGATACACCAAGGAAAGTCTACACGAACCCTTTGGCAAAATCCTGTATATCGTGCGAAAAAGGATGGATATACCGAAAAAATCGCTATAATGACCCCGAAGCAGGGTTATGCAGCGGAAAAGATCCGTCCATGACCAAAATCCCTTAACGTGAGTTTTCGTTCCACTGAGCGTCAGACCCCGTAGAAAAGATCAAAGGATCTTCTTGAGATCCTTTTTTTCTGCGCGTAATCTGCTGCTTGCAAACAAAAAAACCACCGCTACCAGCGGTGGTTTGTTTGCCGGATCAAGAGCTACCAACTCTTTTTCCGAAGGTAACTGGCTTCAGCAGAGCGCAGATACCAAATACTGTTCTTCTAGTGTAGCCGTAGTTAGGCCACCACTTCAAGAACTCTGTAGCACCGCCTACATACCTCGCTCTGCTAATCCTGTTACCAGTGGCTGCTGCCAGTGGCGATAAGTCGTGTCTTACCGGGTTGGACTCAAGACGATAGTTACCGGATAAGGCGCAGCGGTCGGGCTGAACGGGGGGTTCGTGCACACAGCCCAGCTTGGAGCGAACGACCTACACCGAACTGAGATACCTACAGCGTGAGCTATGAGAAAGCGCCACGCTTCCCGAAGGGAGAAAGGCGGACAGGTATCCGGTAAGCGGCAGGGTCGGAACAGGAAGAGCGCACGAGGGAGCTTCCAGGGGGAAACGCCTGGTATCTTTATAGTCCTGTCGGGTTTCGCCACCTCTGACTTGAGCGTCGATTTTTGTGATGCTCGTCAGGGGGGCGGAGCCTATGGAAAAACGCCAGCAACGCGGCCTTTTTACGGTTCCTGGCCTTTTGCTGGCCTTTTGCTCACATGTAATGTGAGTTAGCTCACTCATTAGGCACCCCAGGCTTTACACTTTATGCTTCCGGCTCGTATGTTGTGTGGAATTGTGAGCGGATAACAATTTCACACAGGAAACAGCTATGACCATGATTACGCCAAGCTACGTAATACGACTCACTAGTGGGGCCCgcgccactcggtcgcagggggtgtgggattttttttgggagacaatcctcatggcctatacgacggcccagttggtgactgcgtacaccaacgccaacctcggcaaggcgcctgacgccgccaccacgctgacgctcgacgcgtacgcgactcaaacccagacgggcggcctctcggacgccgctgcgctgaccaacaccctgaagctggtcaacagcacgacggctgttgccatccagacctaccagttcttcaccggcgttgccccgtcggccgctggtctggacttcctggtcgactcgaccaccaacaccaacgacctgaacgacgcgtactactcgaagttcgctcaggaaaaccgcttcatcaacttctcgatcaacctggccacgggcgccggcgccggcgcgacggctttcgccgccgcctacacgggcgtttcgtacgcccagacggtcgccaccgcctatgacaagatcatcggcaacgccgtcgcgaccgccgctggcgtcgacgtcgcggccgccgtggctttcctgagccgccaggccaacatcgactacctgaccgccttcgtgcgcgccaacacgccgttcacggccgctgccgacatcgatctggccgtcaaggccgccctgatcggcaccatcctgaacgccgccacggtgtcgggcatcggtggttacgcgaccgccacggccgcgatgatcaacgacctgtcggacggcgccctgtcgaccgacaacgcggctggcgtgaacctgttcaccgcctatccgtcgtcgggcgtgtcgggttcgGGCGGTAGCGGAGGAGGCTCGGGTGACTACAAGGACGACGATGACAAGGTCCCAGGAGTTGGAGTCCCAGGAGGGGGCGTTCCGGGCGCAGGAGTTCCTGGAGTAGGAGTTCCAGGAGTGGGCGTGCCAGGGGTGGGCGTCCCAGGTGGGGGAGTTCCCGGAGCAGGTGTGCCTGGGGGCGGCGTGCCTGGAGTCGGAGTTCCGGGGGTGGGTGTACCGGGTGGAGGCGTACCAGGCGCGGGAGTGCCGGGCGTGGGCGTGCCAGGCGTCGGTGTACCCGGCGTTGGTGTTCCGGGCGGAGGTGTCCCCGGAGCTGGGGTTCCCGGTGGGGGTGTACCGGGCGTCGGGGTTCCCGGTGTGGGTGTCCCAGGTGGCGGCGTTCCCGGGGCGGGCGTACCTGGAGTGGGTGTGCCAGGAGTCGGCGTCCCAGGAGTCGGCGTACCAGGAGGTGGTGTTCCCGGGGCCGGAGTTCCCGGCGGAGGAGTTCCCGGCGTCGGCGTCCCTGGGGTCGGCGTCCCGGGAGGTGGAGTACCCGGAGCAGGAGTGCCGGGAGTCGGTGTACCTGGTGTCGGTGTCCCTGGTGTAGGTGTCCCGGGTGGTGGGGTGCCAGGTGCTGGCGTACCTGGGGGGGGGGTTCCTGGCGTAGGCGTTCCGGGGGTGGGCGTTCCGGGCGGCGGGGTGCCGGGAGCAGGTGTCCCCGGCGTTGGTGTACCGGGGGTTGGTGTCCCAGGCGTAGGTGTGCCCGGTGGAGGGGTGCCGGGAGCTGGAGTGCCTGGAGGGGGTGTACCAGGGGTCGGTGTTCCCGGTGTAGGAGTACCGGGGGGCGGAGTCCCAGGAGCCGGCGTGCCGGGTGTTGGAGTCCCGGGAGTCGGAGTCCCTGGGGTAGGCGTTCCAGGGGGAGGGGTCCCCGGTGCAGGGGTTCCTGGCGGTGGTGTCCCAGGAGTTGGCGTCCCAGGAGTTGGAGTCCCAGGAGGGGGCGTTCCGGGCGCAGGAGTTCCTGGAGTAGGAGTTCCAGGAGTGGGCGTGCCAGGGGTGGGCGTCCCAGGTGGGGGAGTTCCCGGAGCAGGTGTGCCTGGGGGCGGCGTGCCTGGAGTCGGAGTTCCGGGGGTGGGTGTACCGGGTGGAGGCGTACCAGGCGCGGGAGTGCCGGGCGTGGGCGTGCCAGGCGTCGGTGTACCCGGCGTTGGTGTTCCGGGCGGAGGTGTCCCCGGAGCTGGGGTTCCCGGTGGGGGTGTACCGGGCGTCGGGGGCGGCTCGGGAGGAGGCTCGGGTGCGCATATCGTAATGGTCGATGCATACAAGCCCACGAAAGGAGGTTCAGGCGGCGGAAGCGGTGGTGGAAGCGGAGGTGGGTCAGGCGGAGGCTCAGGGGGAGGTTCGGGTGGCGGGTCTGGAGGAGGCTCGGGTGCTGACCCGGCCTTCGGCGGCTTCGAAACCCTCCGCGTCGCTGGCGCGGCGGCTCAAGGCTCGCACAACGCCAACGGCTTCACGGCTCTGCAACTGGGCGCGACGGCGGGTGCGACGACCTTCACCAACGTTGCGGTGAATGTCGGCCTGACCGTTCTGGCGGCTCCGACCGGTACGACGACCGTGACCCTGGCCAACGCCACGGGCACCTCGGACGTGTTCAACCTGACCCTGTCGTCCTCGGCCGCTCTGGCCGCTGGTACGGTTGCGCTGGCTGGCGTCGAGACGGTGAACATCGCCGCCACCGACACCAACACGACCGCTCACGTCGACACGCTGACGCTGCAAGCCACCTCGGCCAAGTCGATCGTGGTGACGGGCAACGCCGGTCTGAACCTGACCAACACCGGCAACACGGCTGTCACCAGCTTCGACGCCAGCGCCGTCACCGGCACGGGCTCGGCTGTGACCTTCGTGTCGGCCAACACCACGGTGGGTGAAGTCGTCACGATCCGCGGCGGCGCTGGCGCCGACTCGCTGACCGGTTCGGCCACCGCCAATGACACCATCATCGGTGGCGCTGGCGCTGACACCCTGGTCTACACCGGCGGTACGGACACCTTCACGGGTGGCACGGGCGCGGATATCTTCGATATCAACGCTATCGGCACCTCGACCGCTTTCGTGACGATCACCGACGCCGCTGTCGGCGACAAGCTCGACCTCGTCGGCATCTCGACGAACGGCGCTATCGCTGACGGCGCCTTCGGCGCTGCGGTCACCCTGGGCGCTGCTGCGACGCTAGCTTCGGCCGTGACGCGTCTCCGGATGTACAGGCATGCGTCGACCCTCTAGTCAAGGCCTTAAGTGAGTCGTATTACGGACTGGCCGTCGTTTTACAACGTCGTGACTGGGAAAACCCTGGCGTTACCCAACTTAATCGCCTTGCAGCACATCCCCCTTTCGCCAGCTGGCGTAATAGCGAAGAGGCCCGCACCGATCGCCCTTCCCAACAGTTGCGCAGCCTGAATGGCGAATGGCGCTTCGCTTGGTAATAAAGCCCGCTTCGGCGGGCTTTTTTTT |
